# Emergence of long- and short-range functional connectivity parallels the development of neonatal brain gradients

**DOI:** 10.64898/2026.05.19.726280

**Authors:** Qianwen Chang, Stuart Oldham, Sunniva Fenn-Moltu, Nina Treder, Tomoki Arichi, Grainne McAlonan, Dafnis Batalle

## Abstract

The emergence of macroscale brain organisation during the perinatal period represents a critical but poorly understood process. Using resting-state fMRI in 325 term-born neonates within the first month of postnatal life, we characterised the early emergence of macroscale brain organisation with functional gradients. Neonatal functional gradients showed a distinct pattern compared with the canonical adult gradients: the principal gradient in neonates, separating the primary sensorimotor and visual networks, broadly corresponded to the secondary gradient in adults (sensorimotor-visual axis), whereas the secondary gradient in neonates, spanning from primary visual to association networks, aligned with the principal gradient in adults (sensorimotor-association axis). We found that the perinatal emergence of distance-dependent functional connectivity paralleled the development of neonatal functional gradients, potentially underlying the protracted development of the sensorimotor-association hierarchy in neonates. Regions with a higher proportion of short-range functional connectivity were more strongly associated with the sensorimotor-visual gradient in neonates, while regions with a greater proportion of middle- to long-range functional connectivity were more strongly associated with the sensorimotor-association hierarchy. Together, these findings highlight the critical role of short- and long-range functional connectivity, which showed a close correspondence with neonatal macroscale functional hierarchy. Disruptions to this process during the perinatal period may alter macroscale brain organisation and contribute to the early origins of neurodevelopmental conditions.

**Significance Statement:** Macroscale brain organisation, characterised by functional gradients, is a fundamental feature of the adult brain and is closely linked to neurodevelopmental conditions. Characterising its emergence in early life and the underlying mechanisms is therefore critical for understanding typical and atypical brain development. Here, we showed that macroscale brain organisation is present at birth, with neonatal functional gradients showing a broadly reordered pattern relative to canonical adult gradients, highlighting the early maturation of primary visual and sensorimotor networks. We further demonstrated that the perinatal development of long- and short-range functional connectivity parallels the development of neonatal functional gradients, identifying a potential mechanism through which early disruptions in functional connectivity may contribute to later neurodevelopmental outcomes.

## Introduction

Functional gradients summarise high-dimensional functional connectivity into low-dimensional maps, reflecting dominant patterns of variation in functional brain connectivity^1^. In adults, the principal gradient, which captures the largest variation axis in functional connectivity, spans from the association (transmodal) networks to the primary (unimodal) sensorimotor and visual networks. The secondary gradient spans from the sensorimotor network to the visual network^1^. A reordering of these gradients appears to occur from childhood to adolescence^2,3^, with children around 4.5 years of age having a principal gradient following the sensorimotor-visual axis, and the secondary gradient following the sensorimotor-association axis^4^. Around 12-14 years of age, the sensorimotor-association axis then transitions into the principal gradient, while the sensorimotor-visual axis transitions into the secondary gradient, resulting in an adult-like macroscale organisation^2^. While well-characterised in childhood, adolescence, and adulthood, the emergence and development of functional gradients in neonates is less understood. Some studies have suggested that the sensorimotor-association axis is not fully mature at birth, consistent with findings that the association networks are less developed perinatally^5–8^. Among the few existing studies assessing neonatal functional gradients^9–12^, a principal sensorimotor-visual axis, as well as an additional anterior-posterior axis, have been identified. Recent work mapping the trajectories of functional gradients across the lifespan has suggested that the sensorimotor-association gradient begins to emerge during the perinatal period^13^. The protracted development of the sensorimotor-association axis may make it particularly sensitive to a wide range of developmental adversities^14^, with atypical development of this axis observed in individuals with neurodevelopmental conditions^15–17^.

Long-range connections play an important role in brain organisation by integrating distributed brain regions^18–20^. In adults, the organisation of long-range connections has been shown to be aligned with the principal functional gradient (sensorimotor-association axis)^21,22^. Specifically, long-range connections preferentially link the transmodal association regions, whereas sensorimotor regions have relatively more short-range connections^21–23^. Association networks are not fully matured in the neonatal brain and thus exhibit spatially localised connectivity with limited long-range connections, whilst the primary sensorimotor and visual networks show more adult-like network organisation^24^. In keeping with this, prenatal development of functional connections has been found to be primarily limited to short- to middle-range connections^16^, while long-range functional connections are less developed before birth and undergo prolonged maturation throughout the first years of life^25–27^. Evidence suggests that the establishment of a balance between long- and short-range connections is critical during early development; an imbalance of long- and short-range functional connections has been found in neonates and infants with a family history of autism^28,29^. This imbalance can persist into later life^15,30,31^ and is associated with disrupted macroscale brain organisation, reflected in alterations of the sensorimotor-association axis^17^.

In this study, we examined the relationship between the perinatal development of long- and short-range functional connectivity and functional gradients in neonates. We hypothesised: 1) that macroscale brain organisation would emerge early in neonates and resemble patterns observed in early childhood, i.e., in reverse order to adults, with the principal gradient spanning the sensorimotor and visual networks and the secondary gradient spanning the sensorimotor to association axis; and 2) that the emergence of long- and short-range functional connections would be associated with the development of functional gradients. To test this, we first used the developing Human Connectome Project (dHCP) data to characterise functional connectivity gradients in neonates^1^. We then used publicly available adult gradient data to examine the correspondence between functional gradients in neonates and adults across the whole brain, as well as at the system (primary and association regions) and sub-network levels. Finally, we estimated connection length using a novel method based on a precise white matter distance metric, defined as the shortest physical white matter pathway available for each pair of functional connections. We categorised functional connections into long-, middle-, and short-range bins based on connection length and characterised the development of distance-dependent functional connections. Finally, we examined associations between distance-dependent functional connections and functional gradients in neonates across the whole brain, as well as at the system and sub-network levels. We further validated our findings using continuous connectivity distance rather than discrete distance bins, demonstrating the robustness of our findings.

## Results

### Functional gradients in neonates

Group-level functional gradients were characterised in 325 healthy term-born neonates from the dHCP cohort^32^, with gestational age (GA) at birth ≥ 37 weeks, postmenstrual age (PMA) at scan = 41.6 ± 1.67 weeks, and postnatal age (PNA) at scan = 1.56 ± 1.34 weeks. A whole-brain atlas consisting of 400 cortical regions from the Schaefer-400 atlas^33^, 12 subcortical regions from the dHCP template^34^, and 21 cerebellar regions from the SUIT atlas^35^ was created for the neonates in the dHCP 40-week template space^34^. Cortical regions were further assigned to the Yeo-7 networks^36^, and the subcortical and cerebellar regions were treated as separate networks, resulting in 9 sub-networks in total (Figure 1a). Functional connectivity between each pair of regions was calculated as the pairwise Pearson correlation. Functional gradients were then calculated following Margulies et al^1^. The first two gradients, which explained the majority of variance in functional connectivity (41.0% and 14.4%), were selected (Figure 1b). The first gradient (G1_neo_) is anchored at the sensorimotor network at one end and the default-mode network at the other, with the visual network situated toward the end opposite to the sensorimotor network. The second gradient (G2_neo_) is anchored at the primary visual network at one end and association (transmodal) networks at the other (Figure 1c).

**Figure 1.**
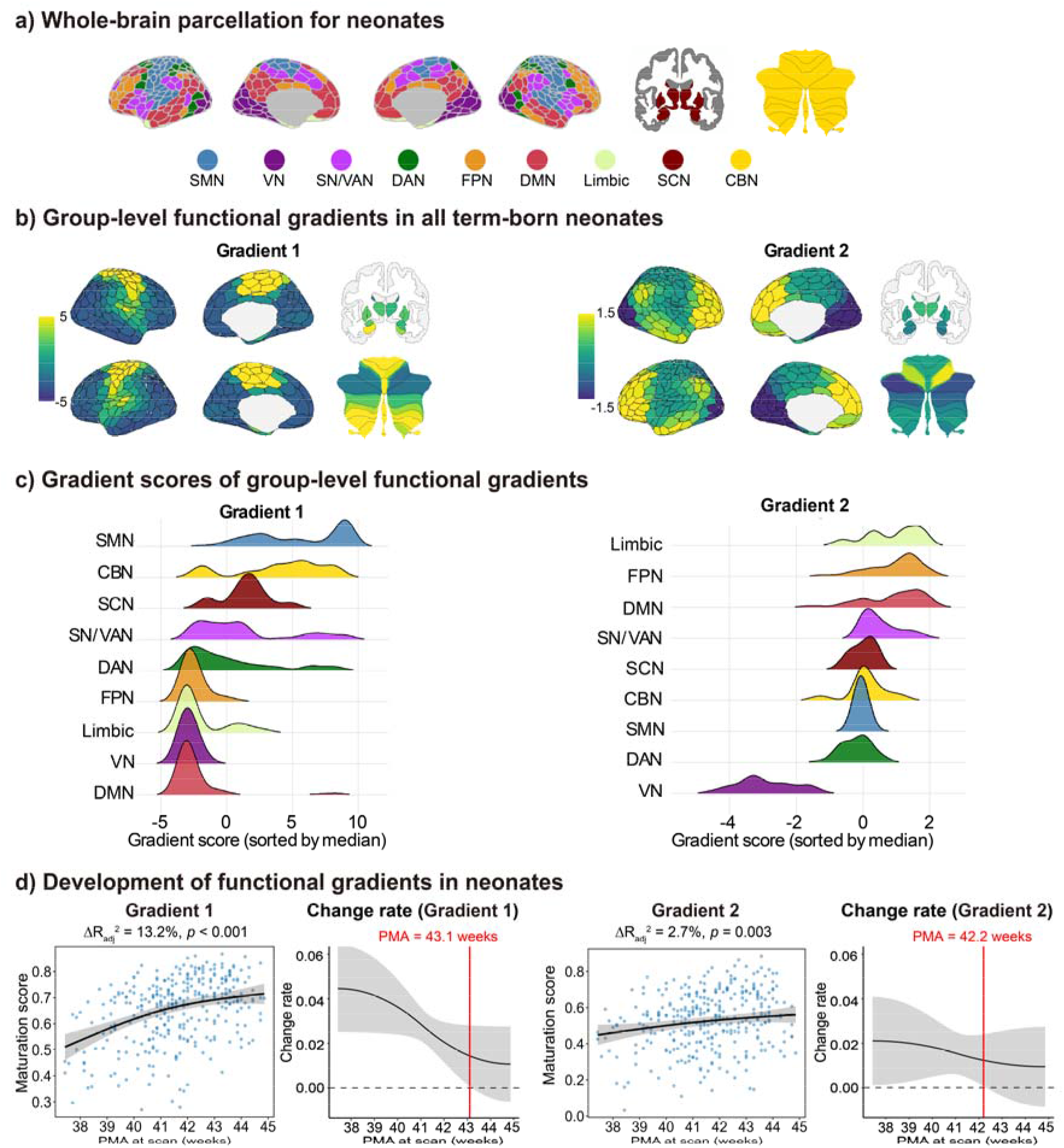
(a) Whole-brain parcellation for neonates. 400 cortical regions from the Schaefer-400 atlas^33^, 12 subcortical regions from the dHCP template^34^, and 21 cerebellar regions from the SUIT atlas^35^ in the dHCP 40-week template space^34^. Cortical regions were assigned to the Yeo-7 networks^36^, and subcortical and cerebellar regions were treated as separate networks, resulting in 9 sub-networks in total. (b) The first two group-level functional gradients in all term-born neonates (325 neonates, GA at birth ≥ 37 weeks). (c) Gradient scores for each sub-network. Sub-networks are ordered by the median gradient score. SMN = sensorimotor network, CBN = cerebellar network, SCN = subcortical network, SN/VAN = salience/ventral attention network, DAN = dorsal attention network, FPN = frontoparietal network, VN = visual network, DMN = default-mode network. (d) Development of functional gradients. Maturation scores of the first two functional gradients significantly increased with PMA at scan, after controlling for sex and in-scanner head motion (FD outliers), modelled using the generalised additive model (GAM). The change rate of maturation score was quantified as the first derivative of the smooth term of PMA at scan.

Individual functional gradients were calculated for each neonate within the same cohort. To examine their development, maturation scores, defined as the Spearman’s rank correlation between individual functional gradients and the group-level functional gradients derived from all term-born neonates, were modelled using a generalised additive model (GAM), with PMA at scan as the smooth term, and sex and in-scanner head motion (framewise displacement [FD] outliers) as linear covariates. Effect size and significance were determined by comparing the full model with a reduced model excluding the smooth term of PMA at scan. We found significant developmental changes in maturation scores for the first two gradients (ΔR_adj_^2^ = 13.2%, *p* < 0.001 for G1_neo_, ΔR_adj_^2^ = 2.7%, *p* = 0.003 for G2_neo_; Figure 1d). For both G1_neo_ and G2_neo_, the change rate of maturation score, measured by the first derivative of the smooth term of PMA at scan, was significantly positive at the lower range of PMA at scan, indicating increasing similarity to the average neonate. The change rate gradually decreased with PMA at scan. For G1_neo_, the first derivative remained significantly positive until approximately 43.1 postmenstrual weeks, after which its 95% confidence interval included zero. For G2_neo_, the corresponding transition occurred at approximately 42.2 postmenstrual weeks. To further examine *in utero* and *ex utero* maturation, we decomposed effects of PMA at scan into GA at birth and PNA at scan. For G1_neo_, we found significant associations with both GA at birth (*r* = 0.411, *p* < 0.001) and PNA at scan (*r* = 0.180, *p* = 0.001). For G2_neo_, we found a significant association with GA at birth (*r* = 0.267, *p* < 0.001), but not with PNA at scan (*r* = 0.015, *p* =0.792).

It should be noted that individual functional gradients were aligned to the group template derived from all the term-born neonates. Therefore, maturation score reflects similarity to the sample average and thus could potentially be influenced by the sampling distribution. As an additional sensitivity analysis, we constructed a reference template using only the oldest term-born neonates (23 neonates, PMA at scan ≥ 44 weeks) and aligned the remaining neonates to this template (Supplementary Figures S1-4). Maturation scores increased significantly with PMA at scan for both the sensorimotor-visual axis (G1; ΔR_adj_^2^ = 14.93%, *p* < 0.001) and the visual-association axis (G3; ΔR_adj_^2^ = 4.70%, *p* < 0.001). Of note, the second gradient extended between the dorsal attention network and the salience/ventral attention network and showed no significant changes (ΔR_adj_^2^ = 0.81%, *p* = 0.098). This sensitivity analysis thus demonstrates that the observed developmental changes cannot be fully explained by sample distribution and alignment target.

### Correspondence between adult and neonatal functional gradients

We next compared the characterised neonatal functional gradients with the canonical adult gradients originally calculated by Margulies et al^1^. The second gradient in neonates (G2_neo_) showed a significant but moderate correspondence with the principal gradient in adults (G1_adult_, sensorimotor-association axis; Spearman’s *r* = 0.626, *p*_spin_ < 0.001, 1000 spin tests; Figure 2a). The first gradient in neonates (G1_neo_) showed a significant but moderate correspondence with the second gradient in adults (G2_adult_, visual-sensorimotor axis; Spearman’s *r* = 0.588, *p*_spin_ = 0.003; Figure 2b).

**Figure 2.**
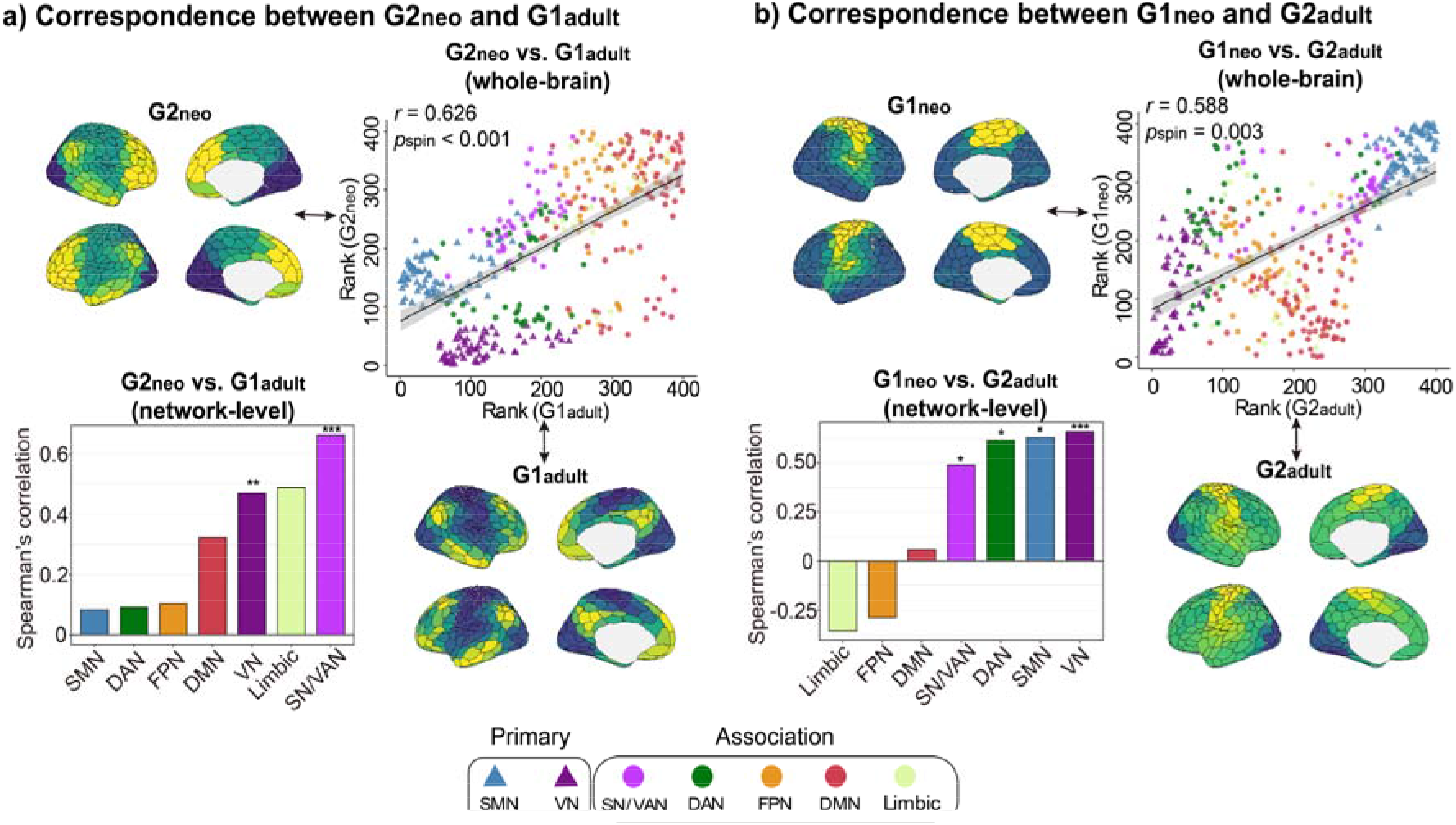
(a) Correspondence between the second gradient in neonates (G2_neo_) and the primary gradient in adults (G1_adult_) at the whole-brain and sub-network levels. (b) Correspondence between the first gradient in neonates (G1_neo_) and the secondary gradient in adults (G2_adult_) at the whole-brain and sub-network levels. Spearman’s rank correlations were calculated between the two brain maps with 1000 spin tests performed for both whole-brain and network level analyses. Adult functional gradient data (cortex-only) were calculated by Margulies et al^1^. SMN = sensorimotor network, SN/VAN = salience/ventral attention network, DAN = dorsal attention network, FPN = frontoparietal network, VN = visual network, DMN = default-mode network. \*\*\**p*_spin_ < 0.001, \*\**p*_spin_ < 0.01, \**p*_spin_ < 0.05.

The Yeo-7 cortical sub-networks were further grouped into primary unimodal (visual and sensorimotor) and association (default mode, frontoparietal, dorsal attention, salience/ventral attention, and limbic) systems. Correspondence between G2_neo_ and G1_adult_ was examined separately within these two systems: significant positive correspondence was found within association regions (Spearman’s *r* = 0.508, *p*_spin_ = 0.005), while no significant correspondence was found within primary unimodal regions (Spearman’s *r* = -0.416, *p*_spin_ = 0.149, 1000 spin tests; Supplementary Figure S5). At the sub-network level, significant correspondence was found in the salience/ventral attention (*r* = 0.662, *p*_spin_ = 0.001, *p*_FDR_ = 0.007, FDR corrected across 7 cortical sub-networks) and visual network (*r* = 0.470, *p*_spin_ = 0.008, *p*_FDR_ = 0.026). Correspondence between G1_neo_ and G2_adult_ was found within primary unimodal regions (Spearman’s *r* = 0.906, *p*_spin_ < 0.001), while no correspondence was found within association regions (Spearman’s *r* = 0.135, *p*_spin_ = 0.251; Supplementary Figure S5). At the sub-network level, significant correspondence was found in the visual, sensorimotor, dorsal attention, and salience/ventral attention networks (*p*_spin_ values < 0.05, Figure 2b). However, after multiple comparison corrections, only the visual network remained statistically significant (Spearman’s *r* = 0.659, *p*_FDR_ < 0.001).

As adult functional gradients were derived from cortico-cortical functional connectivity (i.e. not including subcortical and cerebellar regions), we performed a validation analysis using neonatal functional gradients derived from cortico-cortical functional connectivity only (Supplementary Figure S6). The cortex-only neonatal functional gradients (G1_neo,cort_ and G2_neo,cort_) closely resembled those derived from whole-brain functional connectivity (Spearman’s *r* = 0.990, *p*_spin_ <0.001 for G1_neo_ and G1_neo,cort_; Spearman’s *r* = 0.954, *p*_spin_ < 0.001 for G2_neo_ and G2_neo,cort_) and showed consistent correspondence with adult functional gradients. Specifically, G2_neo,cort_ showed significant correspondence with G1_adult_ (Spearman’s *r* = 0.555, *p*_spin_ < 0.001), and G1_neo,cort_ showed significant correspondence with G2_adult_ (Spearman’s *r* = 0.602, *p*_spin_ = 0.001).

### Development of distance-dependent functional connectivity

Connection lengths, estimated using white matter distance, ranged from 4.10 mm to 122.04 mm. White matter forms the anatomical pathways linking brain regions, providing the structural basis for functional connections. Unlike the widely used Euclidean distance, our white matter distance metric accounts for brain geometry, indicating the shortest route through the interior white matter volume between a pair of regions. Functional connections were categorised into three bins: short-range (<40 mm), middle-range (40-80 mm), and long-range (>80 mm) connections. The proportional contribution of functional connections within each distance bin was quantified using normalised functional connectivity strength (normalised FCS), defined as the sum of connection strengths within a given distance bin divided by a node’s total connection strength. For each node, normalised FCS_long_, normalised FCS_middle_, and normalised FCS_short_ sum to 1.

We used linear regression to examine the development of proportional contributions of short-, middle-, and long-range functional connections, with the PMA at scan as the main predictor, controlling for sex and in-scanner head motion (FD outliers). Effect sizes are reported as the partial correlation coefficient. Long-range functional connections accounted for a greater proportion of total connectivity strength across 7 out of 9 networks with increasing PMA at scan, with significant increases seen in the DMN and frontoparietal networks (FPN) (*r* = 0.167, *p*_FDR_ = 0.038 in the DMN, *r* = 0.146, *p*_FDR_ = 0.024 in the FPN; Figure 3). With increasing PMA at scan, short-range connections accounted for a smaller proportion of total strength in the visual network (*r* = -0.156, *p*_FDR_ = 0.045; Figure 3). No significant changes were observed for middle-range connections within any sub-network (*p*_FDR_ values > 0.05). Despite motion correction and adjustment for % FD outliers, residual head motion may still partly contribute to the observed developmental changes in distance-dependent functional connectivity^37,38^. We therefore repeated the main analyses while additionally controlling for % FD outliers, mean FD, and mean DVARS as a sensitivity analysis. The overall developmental pattern was preserved: long-range functional connections accounted for a greater proportion of total connectivity strength across seven of nine networks with increasing PMA at scan, with nominally significant increases in the frontoparietal and default-mode networks. Conversely, short-range connections accounted for a smaller proportion of total strength with increasing PMA at scan in the visual, frontoparietal, and cerebellar networks, although these network-level effects did not survive FDR correction (Supplementary Figure S7).

**Figure 3.**
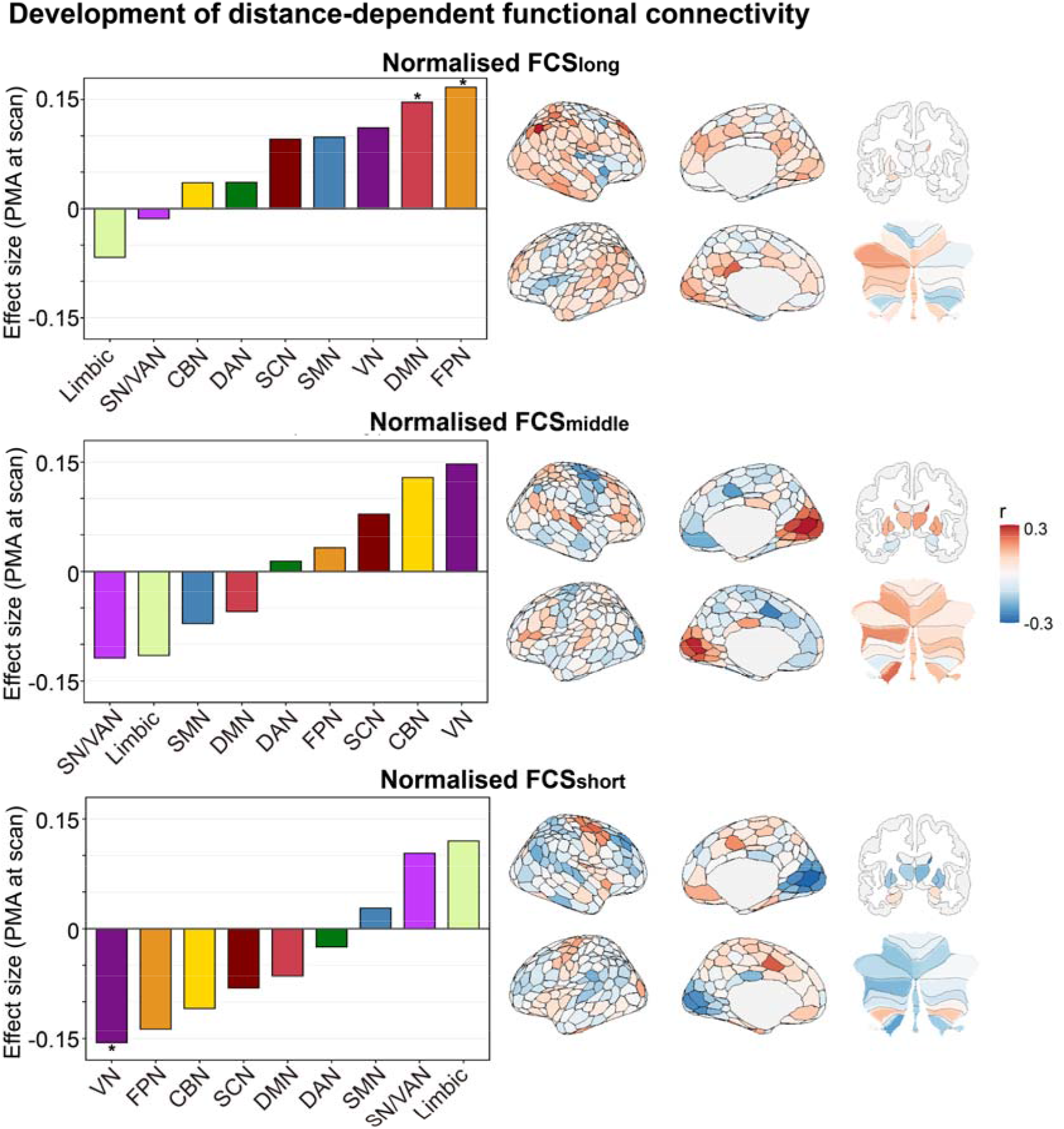
(a) Development of normalised FCS for long-, middle-, and short-range connections at the network (left) and parcel levels (right). Linear regression models were fitted, with PMA at scan as the main predictor, controlling for sex and in-scanner head motion (FD outliers). The effect size was reported as the partial correlation coefficient. SN/VAN = salience/ventral attention network; SMN = sensorimotor network; DMN = default-mode network; DAN = dorsal attention network; FPN = frontoparietal network; VN = visual network; SCN = subcortical network; CBN = cerebellar network. \**p*_FDR_ < 0.05, FDR corrected across 9 networks.

To further examine *in utero* and *ex utero* maturation, we decomposed effects of PMA at scan into GA at birth and PNA at scan. With increasing PNA at scan, normalised FCS_long_ increased across all the sub-networks, with significant increases in the frontoparietal, default-mode, sensorimotor, dorsal attention, and visual sub-networks (*p*_FDR_ values < 0.05). Normalised FCS_short_ decreased in the visual, frontoparietal, and dorsal attention sub-networks (*p*_FDR_ values < 0.05). In contrast, associations with GA at birth were more limited, with a significant positive association observed only between GA at birth and normalised FCS_short_ in the limbic network (*r* = 0.154, *p*_FDR_ = 0.04996; Supplementary Figure S8).

### Associations between distance-dependent functional connectivity and functional gradients

To investigate the relationship between long- and short-range functional connectivity and the development of neonatal functional gradients, we examined associations between functional gradient maps and normalised FCS maps at the group level. Spearman’s rank correlations were first evaluated between the group-averaged normalised FCS maps and the group-level gradient templates across the whole brain, followed by system-level analyses of primary and association regions and network-level analyses.

Globally, the G1_neo_ showed distinct associations with long- and short-range functional connections, with a negative association with normalised FCS_long_ (Spearman’s *r* = - 0.532, *p* < 0.001; validated when assessed with 1000 spin tests for cortical regions only, *r* = -0.545, *p*_spin_ = 0.011; Figure 4a (i)), but a positive association with normalised FCS_short_ (Spearman’s *r* = 0.443, *p* < 0.001; with 1000 spin tests for cortical regions, *r*= 0.467, *p*_spin_ = 0.008; Figure 4a (ii)). At the system level, negative associations with normalised FCS_long_ were found within both primary (Spearman’s *r* = -0.558, *p*_spin_ = 0.048) and association regions (Spearman’s *r* = -0.544, *p*_spin_ = 0.006), while positive associations with normalised FCS_short_ were found in association regions (Spearman’s *r* = 0.499, *p*_spin_ = 0.007) but not in primary regions (Spearman’s *r* = 0.307, *p*_spin_ = 0.170). Within cortical sub-networks, significant negative correlations with long-range connections were found in dorsal attention, salience/ventral attention, frontoparietal, and default-mode networks (*p*_spin_ values < 0.05). After multiple comparison correction, these negative correlations remained statistically significant only in the frontoparietal and dorsal attention networks (*p*_spin_ = 0.014, *p*_FDR_ = 0.047 in the FPN, *p*_spin_ = 0.011, *p*_FDR_ = 0.047 in the DAN, Figure 4b (i)). No statistically significant associations were found with short-range connections, though all sub-networks showed a positive correlation (*p*_spin_ values > 0.05; Figure 4b (ii)).

**Figure 4.**
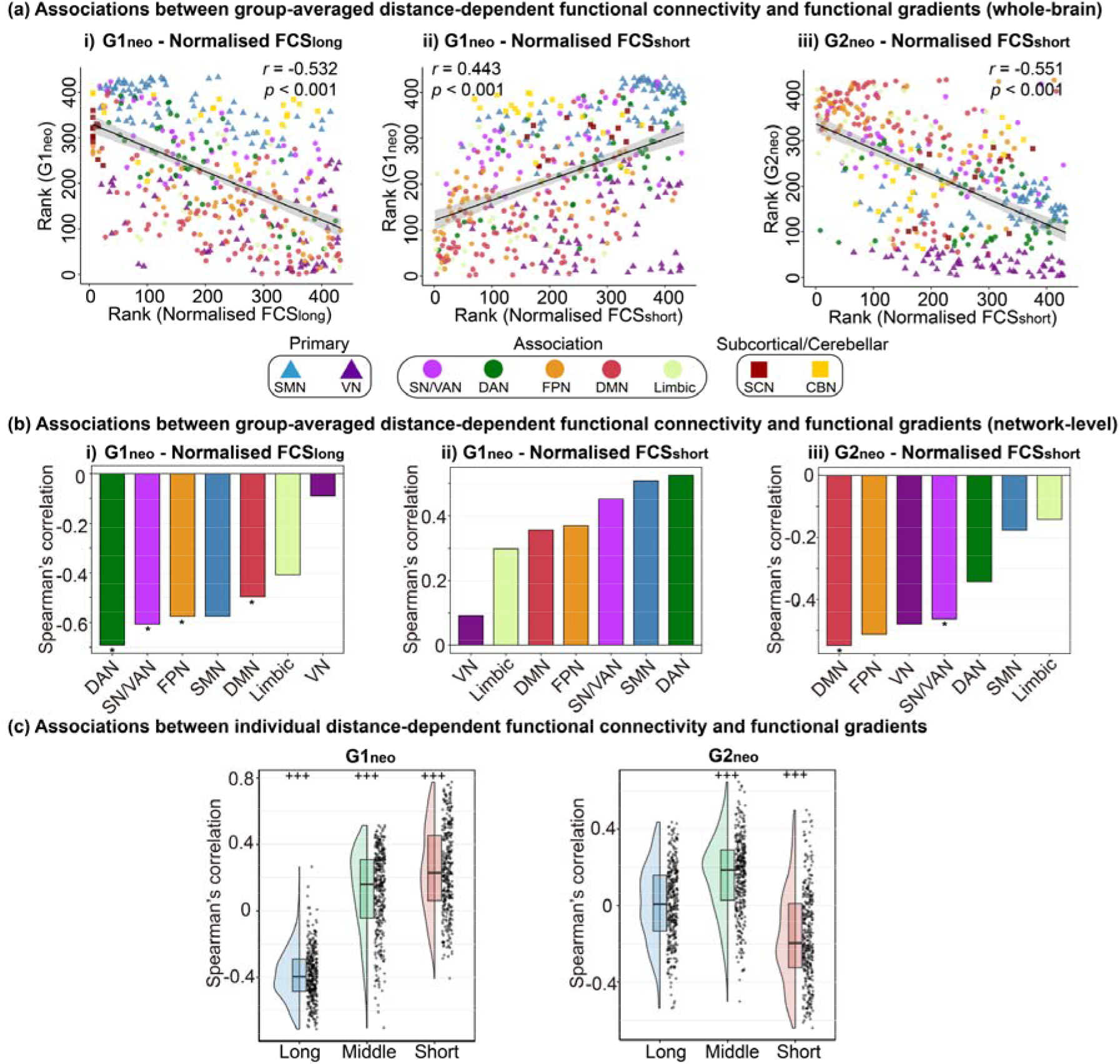
Associations between group-averaged distance-dependent functional connectivity and functional gradients at the (a) whole-brain and (b) network levels. Spearman’s rank correlations were calculated between the group-averaged normalised FCS and the group templates of functional gradient maps. Only associations significant at the whole-brain level after spin tests are shown, and corresponding network-level results are shown for these associations. SN/VAN = salience/ventral attention network; SMN = sensorimotor network; DMN = default-mode network; DAN = dorsal attention network; FPN = frontoparietal network; VN = visual network. \**p*_spin_ < 0.05. Associations between individual distance-dependent functional connectivity and functional gradients across the whole-brain. Spearman’s rank correlations were calculated between normalised FCS (of long, middle and short-range connections) and functional gradient maps for each neonate. ^+++^*p*_FDR_ < 0.001.

The G2_neo_ was negatively associated with normalised FCS_short_ (Spearman’s *r* = -0.551, *p* < 0.001; with 1000 spin tests for cortical regions, *r* = -0.582, *p*_spin_ = 0.002; Figure 4a (iii)). Whilst there were no significant correlations with normalised FCS_middle_ and normalised FCS_long_ individually (*p*_spin_ values > 0.05), negative correlation with short-range connections is equivalent to a positive association with the overall proportion of middle- to long-range connections (i.e., Spearman’s *r* = 0.551, *p* < 0.001; with 1000 spin tests for cortical regions, *r* = 0.582, *p*_spin_ = 0.002), given that the proportional contributions sum to 1. At the system-level, negative association with short-range connections was found within association regions (Spearman’s *r* = -0.563, *p*_spin_ = 0.008), but not in primary regions (Spearman’s *r* = -0.011, *p*_spin_ = 0.494). At the sub-network level, negative correlations were found in the DMN and SN/VAN (*p*_spin_ = 0.040 in the DMN, *p*_spin_ = 0.024 in the SN/VAN; Figure 4b (iii)). However, none of the associations survived multiple comparisons correction (*p*_FDR_ values > 0.05).

To complement the group-level analyses, we further examined associations at the individual level. For each individual, Spearman’s rank correlations were calculated between each normalised FCS map (short-, middle-, and long-range) and each gradient map (G1_neo_ and G2_neo_), resulting in six correlations per individual. One-sample t-tests were then performed to examine whether each correlation was significantly different from zero across neonates (Figure 4c). A similar pattern was observed at the individual level: G1_neo_ was negatively associated with normalised FCS_long_ (Mean ± SD: Spearman’s *r* = -0.384 ± 0.156; *t* = -44.243, *p*_FDR_ < 0.001, FDR corrected for 6 comparisons) and positively associated with both middle-range (Spearman’s *r* = 0.117 ± 0.251; *t* = 8.385, *p*_FDR_ < 0.001) and short-range connections (Spearman’s *r* = 0.245 ± 0.259; *t* = 17.010, *p*_FDR_ < 0.001). In contrast, G2_neo_ was negatively associated with short-range connections (Mean ± SD: Spearman’s *r* = -0.159 ± 0.243; *t* = -11.791, *p*_FDR_ < 0.001), positively associated with middle-range connections (Spearman’s *r* = 0.158 ± 0.201; *t* = 14.142, *p*_FDR_ < 0.001), and showed no significant association with normalised FCS_long_ (Mean ± SD: Spearman’s *r* = 0.008 ± 0.203; *t* = 0.741, *p*_FDR_ = 0.459). We next examined whether these correlations between distance-dependent functional connectivity and gradient maps changed with PMA at scan using linear regression, controlling for sex and in-scanner head motion (Supplementary Figure S9). With increasing PMA at scan, the correlation between G1_neo_ and normalised FCS_long_ decreased (*r* = -0.110, *p*_UNC_ = 0.048), while the correlation between G1_neo_ and normalised FCS_short_ increased (*r* = 0.117, *p*_UNC_ = 0.035), though these correlations did not survive correction for multiple comparisons. No significant developmental changes were found for the remaining correlations (*p* values > 0.05).

As neonatal functional gradients and normalised FCS maps were derived from the same functional connectivity matrix, the two measures are not fully independent. We thus performed an additional validation analysis consisting of a split-half analysis to reduce the dependency. Specifically, we divided each neonate’s resting-state data timeseries into two halves and estimated functional gradients from the first half and distance-dependent functional connectivity from the second half. We then repeated the key analyses using these separately estimated measures (Supplementary Figures S10-13). The relationships between neonatal functional gradients and distance-dependent functional connectivity were replicated, showing that the observed relationships are unlikely to solely driven by estimating both measures from the same connectivity matrix.

### Connectivity distance as a complementary measure to normalised FCS within distance bins

We then computed nodal *connectivity distance*, a continuous measure defined as the average connection length of a node to its functionally connected nodes after proportional thresholding^21^. To complement the normalised FCS results within each distance bin, connectivity distance was examined using the same analytical framework. Connectivity distance in the frontoparietal and visual networks increased significantly with PMA at scan (*r* = 0.152, *p*_FDR_ *=* 0.028 in the FPN*, r =* 0.158, *p*_FDR_ = 0.028 in the visual network; Figure 5a). The G1_neo_ was negatively correlated with the connectivity distance map (Spearman’s *r* = -0.550, *p* < 0.001; with 1000 spin tests in cortical regions, *r* = -0.550, *p*_spin_ = 0.008; Figure 5b (i)), indicating that regions with longer connectivity distances had lower G1_neo_ scores. At the system level, significant negative correlations were found in the association regions (Spearman’s *r* = -0.568, *p*_spin_ = 0.003), but not within the primary unimodal regions (Spearman’s *r* = -0.473, *p*_spin_ = 0.078). Across cortical sub-networks, while correlations were negative, none of these associations survived multiple comparison correction (*p*_FDR_ values > 0.05).

**Figure 5.**
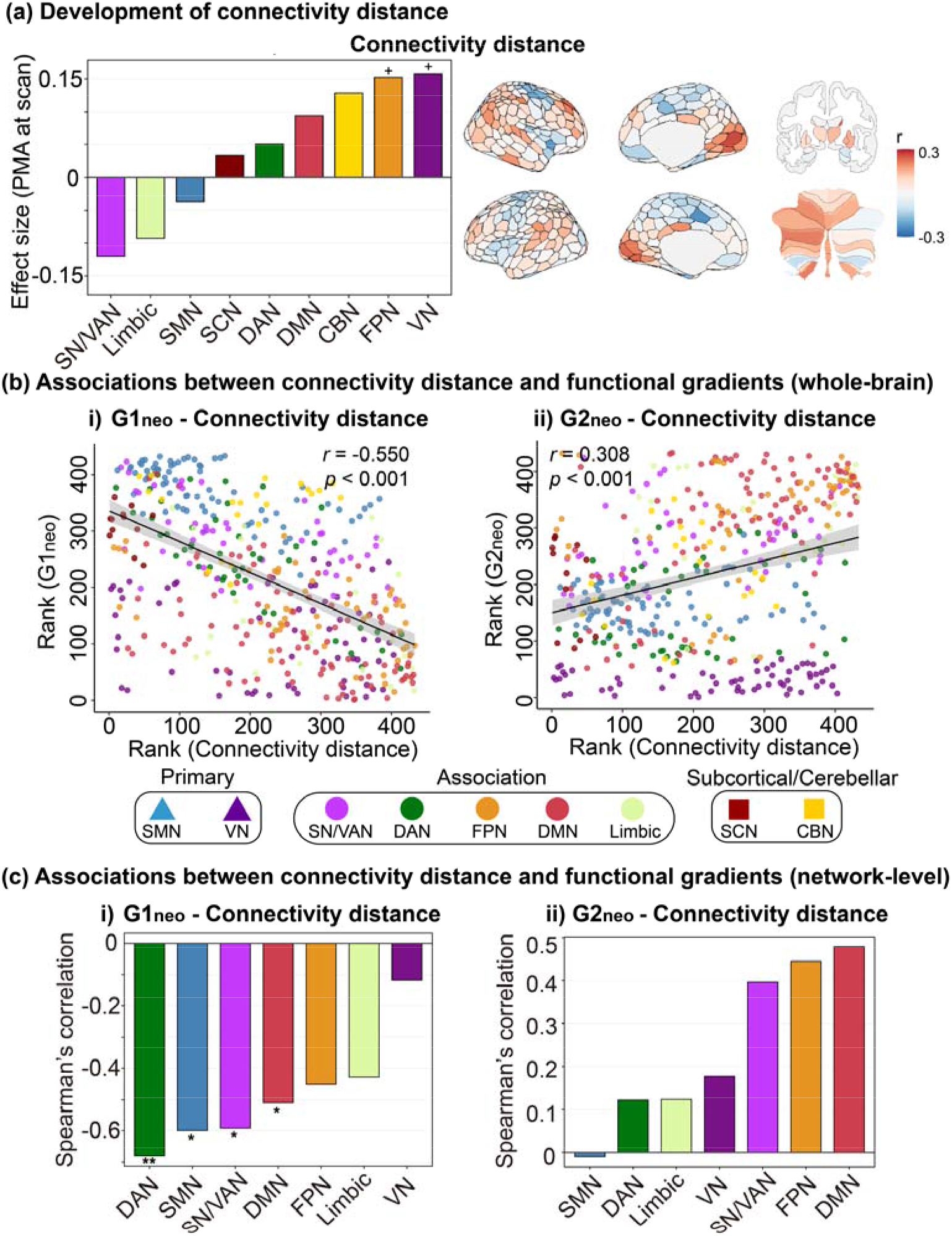
(a) Development of connectivity distance at the network (left) and parcel levels (right). A linear regression model was fitted to connectivity distance, with PMA at scan as the main predictor, controlling for sex and in-scanner head motion (FD outliers). The effect size was reported as the partial correlation coefficient. ^+^*p*_FDR_ < 0.05, FDR corrected across 9 networks. Associations between group-averaged connectivity distance and functional gradients at the (b) whole-brain and (c) network levels. Spearman’s rank correlations were calculated between the group-averaged connectivity distance and the group template of functional gradient maps. SN/VAN = salience/ventral attention network; SMN = sensorimotor network; DMN = default-mode network; DAN = dorsal attention network; FPN = frontoparietal network; VN = visual network; SCN = subcortical network; CBN = cerebellar network. \*\**p*_spin_ < 0.01, \**p*_spin_ < 0.05.

For the G2_neo_, no statistically significant correlation was observed with the connectivity distance map globally (Spearman’s *r* = 0.308, *p* < 0.001; with 1000 spin tests in cortical regions, *r* = 0.318, *p*_spin_ = 0.082; Figure 5b (ii)). At the system level, a significant positive correlation was found within association regions (Spearman’s *r* = 0.441, *p*_spin_ = 0.024), but not within primary unimodal regions (Spearman’s *r* = -0.237, *p*_spin_ = 0.255). No significant correlations were found within any cortical sub-network for G2_neo_ (all *p*_spin_ values > 0.05; Figure 5c (ii)).

## Discussion

In this study, we identified functional gradients in neonates, characterising the early emergence of macroscale brain organisation, and demonstrating how distance-dependent functional connectivity is closely linked with gradient development. The first two functional gradients explained most of the variance in functional connectivity. The first neonatal gradient, anchored at the sensorimotor network on one end and the visual network toward the other end, showed significant correspondence with the secondary gradient in adults (sensorimotor-visual axis). The second neonatal gradient, spanning the association and visual networks, showed significant correspondence with the primary gradient in adults (sensorimotor-association axis). Furthermore, functional gradients changed significantly with PMA at scan: with increasing age, individual functional gradients increasingly resembled those of the average neonate. During early development, the proportion of long-range functional connectivity increased globally, with significant increases observed in the default-mode and frontoparietal networks. We further demonstrated that distance-dependent functional connectivity parallels the early emergence of macroscale brain organisation: regions with more short-range functional connections showed higher gradient scores in the first gradient, while regions with more middle- to long-range functional connections showed higher gradient scores in the second gradient. These findings were robust when a continuous measure of connectivity distance was used.

The first functional gradient in neonates, which explains the largest variance in functional connectivity, separates the primary sensorimotor and visual networks. Previous studies have also identified this principal sensorimotor-visual axis in neonates prior to 37 postmenstrual weeks^9^. This sensorimotor-visual axis is consistent with evidence that primary unimodal cortices are relatively mature and roughly approximate their adult configuration during the perinatal period, while association regions are less developed^5–8^. Functional hubs are also initially located in primary unimodal cortices before gradually shifting toward association regions later in life^39^. In addition, we found that maturation scores in the first gradient increased with PMA at scan. When we decomposed the effects of PMA at scan into GA at birth and PNA at scan, both were significantly associated with maturation score, suggesting that both the *in utero* environment during gestation and postnatal experience contribute to the development of this gradient. Within this gradient, the sensorimotor and visual networks showed greater similarity to the canonical adult gradient than association networks. Together, this may reflect postnatal experience driving the rapid development of primary motor, visual, and auditory functions after birth^40^, which then provide essential input and output for the higher-order functions supported by association networks, thereby establishing a foundation for hierarchical brain development^8^.

We identified a second functional gradient spanning from primary unimodal to association regions, which corresponds to the primary gradient in adults (sensorimotor-association axis). Prior studies reported an anterior-posterior axis as the secondary gradient in neonates, and suggested that it may develop into the sensorimotor-association axis later in life^3,10^. A recent study, examining the lifespan development of the sensorimotor-association hierarchy, identified an anterior-posterior axis before the first month of life^13^. In our study, we observed a more pronounced pattern differentiating primary unimodal and association regions, with higher gradient scores in the default-mode, frontoparietal, and limbic networks (including prefrontal, temporal, and inferior parietal cortices) and lower scores in primary sensorimotor and visual networks. This may reflect methodological differences: we used the Schaefer-400 atlas, while previous studies^3,10^ used voxel- or vertex-wise analyses. In addition, our functional gradient templates were derived from all term-born neonates available, instead of a sample selection in different time windows as done in prior work^9^. However, when using a subgroup of the oldest term-born neonates (23 neonates, PMA at scan ≥ 44 weeks), this axis was also reliably identified and showed a significant and moderate correspondence with the primary gradient in adults (see Supplementary Figures S1-4). This finding is in keeping with earlier work by Larivière et al^10^ using 40 term-born neonates, which found a significant association between the anterior-posterior gradient in neonates and the primary gradient in adults. Similarly, a recent study, which decomposed functional connectivity into functional harmonics, also reported an adult-like sensory-to-transmodal component that significantly associated with the primary functional gradient in adults^12^. This functional gradient may reflect the anterior-posterior neurogenesis process, myelination^41^, and gene expression^42,43^. In our data, gradient maturation scores significantly increased with PMA at scan, consistent with development towards an adult-like organisation. Extending previous studies reporting an adult-like anterior-posterior axis in neonates^10^, we demonstrated its typical development within the first month of postnatal life. Importantly, this development is found in a larger cohort, highlighting the early emergence and rapid refinement of macroscale cortical organisation.

Correspondence between neonatal and adult functional gradients suggests early emergence of macroscale brain organisation, although their ordering is reversed: the sensorimotor-association axis is the principal gradient in adults, while the sensorimotor-visual axis dominates in neonates. A similar pattern has been observed in preschool children at 4.5 years of age^4^, with additional evidence suggesting that the sensorimotor-visual axis remains the primary gradient until late childhood, after which functional gradients reverse order during adolescence^2,3^. This developmental trajectory is consistent with evidence that association regions mature after primary unimodal areas, paralleling the development of cognitive functions from primary sensory processing to higher-level executive functions^44^. Dividing cortical networks into primary and association regions revealed distinct patterns: G1_neo_ positively correlated with G2_adult_ only in primary unimodal regions, indicating that sensorimotor and visual networks are already established and anchored at opposite ends of this gradient at birth. G2_neo_ positively correlated with G1_adult_ in association regions. Although primary sensorimotor and visual networks are both located at the lower end in neonatal G2_neo_ and adult G1_adult_, no significant association was found within the primary regions. Together, these findings suggest that primary unimodal networks establish early, while the sensorimotor-association axis undergoes prolonged maturation. This late maturation of the sensorimotor-association axis parallels the protracted development of long-range functional connections^25,45^ and may make it vulnerable to disruption in neurodevelopmental conditions. Of note, although our findings suggest a broad developmental reordering between neonatal and adult functional gradients, it is not a strict one-to-one mapping (G2_neo_ and G2_adult_: Spearman’s *r* = 0.329, *p*_spin_ = 0.148; G1_neo_ and G1_adult_: Spearman’s *r* = -0.613, *p*_spin_ < 0.001). G1_neo_ was associated with both G2_adult_ and G1_adult_, which may reflect its spatial pattern. In G1_neo_, both the visual network and association networks (i.e., default-mode, limbic, and frontoparietal networks) positioned opposite the sensorimotor network. Consistent with this interpretation, the adult sensorimotor-visual axis reported by Xia et al.^3^, in which the visual and association networks are similarly positioned opposite the sensorimotor network, visually resembled our G1_neo_. Overall, these findings suggest a broad, but not a strict one-to-one, reordering of functional gradients between neonates and adults.

Our results suggest that distance-dependent functional connectivity is associated with early macroscale brain organisation. Regions with predominantly short-range connections showed higher scores in G1_neo_, while regions with more middle- to long-range connections showed higher scores in G2_neo_. We found that the proportion of total connectivity strength attributed to long-range connections increased, with notable increases in the default-mode and frontoparietal networks during early development. The strengthening of long-range functional connections, especially within association networks, likely supports the formation and development of the sensorimotor-association axis, and the protracted development of long-range functional connections likely further contributes to the dominance of the sensorimotor-association axis in adulthood. The relationship between distance-dependent functional connectivity and the sensorimotor-association axis has been shown in adults, where long-range functional connections are distributed along the sensorimotor-association axis^21,22^. Further supporting this relationship, autism studies have linked atypical functional gradients with reduced long-range functional connections, especially in transmodal regions^15,30^. When examining primary unimodal and association regions separately, we observed similar relationships between the first gradient and distance-dependent functional connectivity across the whole brain. For the second gradient in neonates, a similar relationship was found within association regions but not primary unimodal regions. The findings of these categorical analyses were also consistent with the results using a continuous measure of connectivity distance. Together, these findings indicate that the balance between long- and short-range functional connections is linked to the development of early functional gradients. This could also reflect a dynamic interplay where emerging functional connectivity and macroscale brain organisation mutually influence one another during brain maturation. While associations between these measures were demonstrated and replicated using a split-half analysis, the causal mechanisms linking long- and short-range functional connectivity to the development of functional gradients remain unclear.

There are several strengths and limitations in this study. First, associations between distance-dependent functional connectivity and functional gradients were based on group-averaged analyses in all term-born neonates. While we replicated our findings at both the individual level and group level across a subgroup of the oldest term-born neonates, future studies will likely benefit from characterising inter-individual variability across all ages during perinatal development. Second, analyses were performed in the dHCP week-40 template space rather than individual native space. Therefore, we are unable to assess how individual variability in white matter distance may shape functional connectivity. Third, we used white matter distance to estimate connection length. This measure may provide a closer approximation of potential anatomical pathways than Euclidean or geodesic distance because it constrains paths to the white matter. Nevertheless, it should be interpreted as a white-matter-constrained distance measure rather than an estimate of actual tract length, as it does not incorporate information about fibre orientation or tract topology. Without prior reference, we defined long- and short-range connections based on the empirical distribution of functional connection lengths, ensuring only a small proportion qualified as long-range (see Methods). It should be noted that the assessment of long- and short-range functional connectivity near the 40mm and 80mm bin ages may be affected by brain size. To address this, we validated our findings using connectivity distance as a continuous measure, showing consistent results. Although the dHCP preprocessing pipeline uses ICA-FIX to identify and regress out noise components associated with cardiorespiratory fluctuations, it may not fully remove the aliased signals related to higher frequency cardiovascular pulsations and their potential distance-dependent spatial effects^46^. Regardless, these effects are unlikely to explain the observed age-dependent increases in proportional long-range connectivity and/or the network-specific results, as these are unlikely to covary with cardiac pulsation artefacts. Residual head motion has been reported to partly contribute to the observed developmental changes in distance-dependent functional connectivity^37,38^. When we repeated the analysis with mean FD and mean DVARS (in addition to % FD outliers) in order to further account for residual head motion, the principal pattern in the development of distance-dependent functional connectivity was preserved (Supplementary Figure S7). However, the network-level effects were attenuated following this additional adjustment, and residual motion-related effects cannot be excluded completely. Furthermore, we used the Schaefer atlas with the corresponding Yeo-7 network partitions to ensure generalisability and enable comparison with the well-described adult model of functional organisation. While this atlas has been used in neonatal studies^47–50^ and allowed us to examine distinct patterns in primary unimodal and association regions, we acknowledge that an adult atlas may not fully reflect the neonatal brain’s functional organisation. Future studies may thus benefit from using neonate-specific atlases. Lastly, subcortical regions and the cerebellum were excluded from system-level and network-level analyses due to constraints in network assignment and spin tests. Extending these analyses to subcortical and cerebellar structures could further reveal their contributions to early gradient organisation. The cerebellum forms reciprocal connections with the cerebral cortex in adults^51–53^, which emerge early in life^54^. In addition, thalamocortical connections undergo rapid postnatal development and are critical for cortical specialisation in neonates^55–57^.

In conclusion, our study demonstrates the early emergence of macroscale brain organisation at birth. We find that the sensorimotor-visual and primary-association axes are already present and developing at birth, although their order is reversed compared to adults. Functional gradients in neonates fit into the developmental trajectory of macroscale brain organisation from early childhood through adolescence to adulthood, extending this trajectory into the earliest postnatal period. We further show that the perinatal strengthening of long-range functional connections is closely linked with the development of neonatal functional gradients. Specifically, the first gradient separating sensorimotor and visual networks is linked to the distribution of short-range functional connections, while the second gradient separating primary and association networks corresponds to the distribution of middle- to long-range functional connections. During early development, increases in long-range connections, especially within association regions, may support the formation of the sensorimotor-association axis. Together with previous findings linking reduced long-range connectivity and atypical functional gradients in autism, our results suggest that the perinatal period may be a critical window when disruptions in functional connectivity could alter macroscale brain organisation, potentially leading to the early origins of neurodevelopmental conditions.

## Materials and Methods

### Participants

A total of 325 term-born neonates (gestational age at birth ≥ 37 weeks) from the developing Human Connectome Project (dHCP, release 3)^32^ were included in this study. Inclusion and exclusion criteria, as well as distributions of PMA at scan, GA at birth, and PNA at scan, are provided in Figure 6. Resting-state fMRI (rs-fMRI) was acquired at term equivalent age (postmenstrual age ≥ 37 weeks) during natural sleep. Demographic information are shown in Table 1. PMA at scan was not significantly associated with mean FD (*r* = -0.051, *p* = 0.359) but was negatively associated with % FD outliers (*r* = - 0.209, *p* < 0.001) and positively associated with mean DVARS (*r* = 0.179, *p* = 0.001).

**Figure 6.**
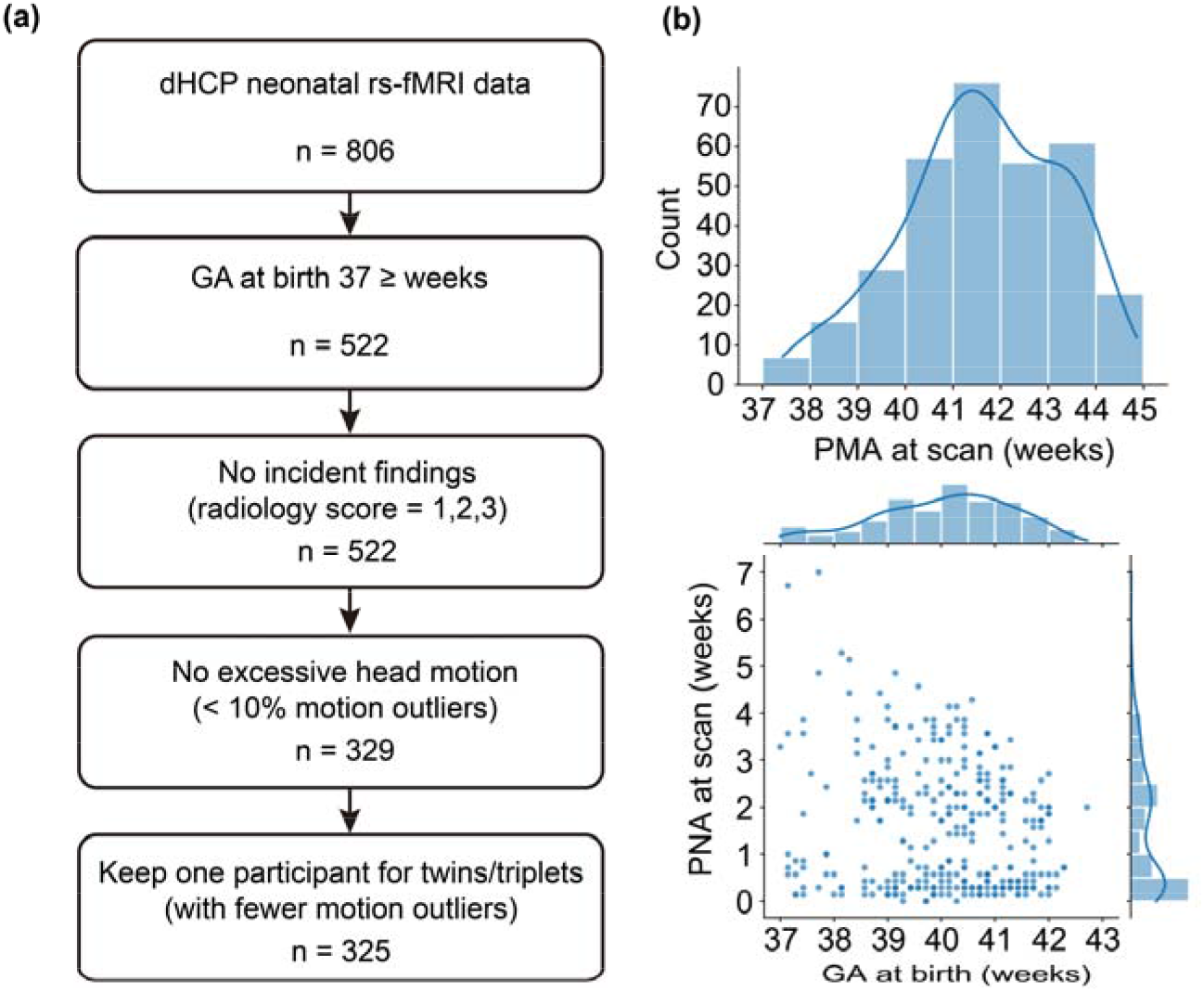
(a) Inclusion and exclusion criteria. (b) Distributions of postmenstrual age (PMA) at scan, gestational age (GA) at birth, and postnatal age (PNA) at scan.

**Table 1.**
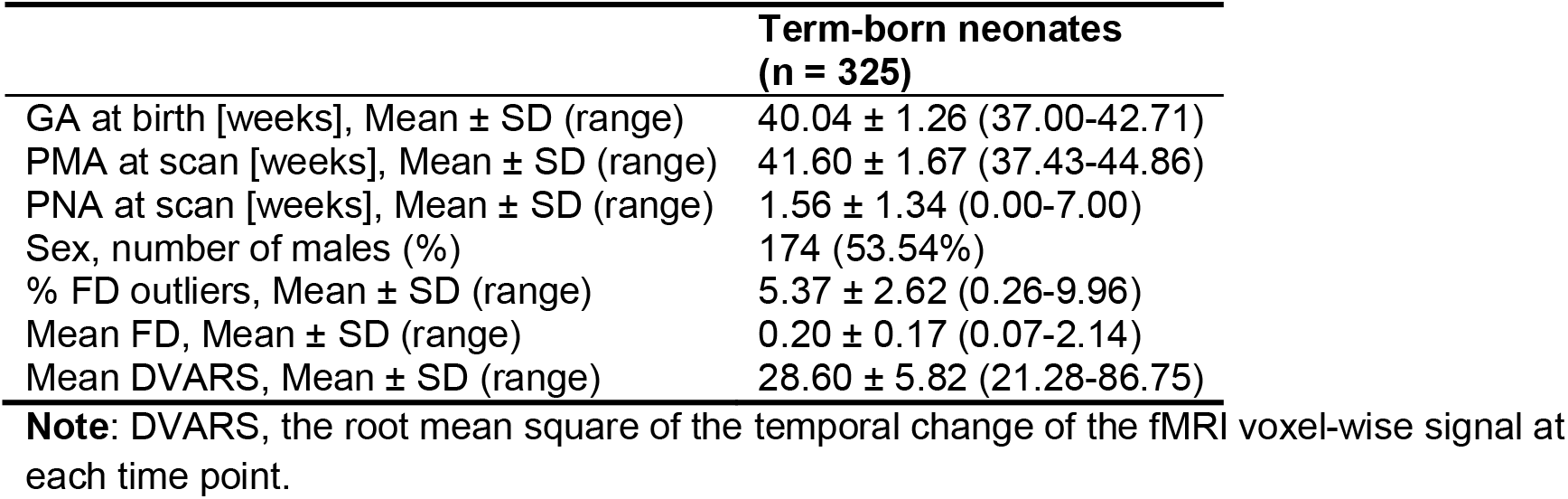
Demographic information.

|  | Term-born neonates<br>(n = 325) |
| --- | --- |
| GA at birth [weeks], Mean ± SD (range) | 40.04 ± 1.26 (37.00-42.71) |
| PMA at scan [weeks], Mean ± SD (range) | 41.60 ± 1.67 (37.43-44.86) |
| PNA at scan [weeks], Mean ± SD (range) | 1.56 ± 1.34 (0.00-7.00) |
| Sex, number of males (%) | 174 (53.54%) |
| % FD outliers, Mean ± SD (range) | 5.37 ± 2.62 (0.26-9.96) |
| Mean FD, Mean ± SD (range) | 0.20 ± 0.17 (0.07-2.14) |
| Mean DVARS, Mean ± SD (range) | 28.60 ± 5.82 (21.28-86.75) |

**Note:** DVARS, the root mean square of the temporal change of the fMRI voxel-wise signal at each time point.

### MRI data acquisition and preprocessing

MRI data were collected at the Evelina Newborn Imaging Centre, Evelina London Children’s Hospital, using a 3 Tesla Philips Achieva system with a 32-channel neonatal head coil^58^. Resting-state fMRI was acquired using a multi-slice echo planar imaging sequence with multiband excitation (factor 9) (repetition time (TR) = 392 ms, echo time (TE) = 38 ms, voxel size = 2.15 × 2.15 × 2.15 mm, flip angle = 34^°^, 45 slices, number of volumes = 2300). T1-weighted images had a reconstructed spatial resolution = 0.8 × 0.8 × 0.8 mm, field of view = 145 ×122 × 100 mm, TR = 4795 ms. T2-weighted images had a reconstructed spatial resolution = 0.8 × 0.8 × 0.8 mm, field of view = 145 × 145 × 108 mm, TR = 12 s, TE = 156 ms^32^.

fMRI data were preprocessed using the dHCP neonatal pipeline^59^, which included dynamic distortion and intra- and inter-volume motion correction using slice-to-volume and rigid-body registration, high-pass filtering with a 150s cutoff. Residual motion, multiband acquisition, and cardiorespiratory artefacts were regressed out with single-subject ICA noise components identified using FSL FIX^60^. Data were subsequently registered to T2w native space using boundary-based registration and then nonlinearly transformed into a 40-week neonatal template from the extended dHCP volumetric atlas^34^. fMRI data with excessive motion (more than 10% volumes classified as motion outliers, i.e., framewise displacement [FD] > 75th centile + 1.5 IQR) or incidental MRI findings of clinical significance (radiology score = 4 or 5) were excluded^61,62^. The data were bandpass filtered at 0.01-0.1Hz^61^.

### Whole-brain atlas

To ensure generalisability, an anatomically defined whole-brain atlas was constructed. The atlas comprises 400 cortical regions from the Schaefer atlas (400 parcels)^33^, which has been used in previous neonatal studies^47–50^. 12 Subcortical regions were taken from the dHCP template^34^, and 21 cerebellar regions (i.e., bilateral lobules I-IV, V, VI, Crus I, Crus II, VIIB, VIIIA, VIIIB, IX, X, and vermis) from the SUIT atlas^35^, making a total of 433 regions. The atlas was nonlinearly transformed to the extended dHCP 40-week template space^34^ using the advanced normalisation tools (ANTs) SyN algorithm^63^. Cortical parcels were assigned to the Yeo-7 sub-networks^36^. Subcortical and cerebellar parcels were treated as separate networks, resulting in 9 sub-networks in total.

### Functional connectivity

Functional connectivity (FC) was computed as the pairwise Pearson’s correlation between the average timeseries of 433 parcels, with only positive correlations retained. This was done as the biological meaning of negative functional connectivity is unknown, and thus its interpretation remains controversial. To calculate group-level FC, individual FC matrices were Fisher r-to-z transformed, averaged across participants, and then transformed back to correlation coefficients using the Fisher z-to-r transform.

### Neonatal functional gradients

Neonatal functional gradients were derived using the approach described by Margulies et al.^1^. A 10% row-wise threshold was applied to the functional connectivity matrix, and cosine similarity was calculated between the functional connectivity profiles of each pair of nodes. Diffusion map embedding was then used to reduce the high-dimensional similarity matrix to low-dimensional gradients. Individual gradients were aligned to a group template using Procrustes rotation, accounting for potential differences in gradient sign and order. The group template was defined from the group-averaged functional connectivity matrix of all term-born neonates. For each neonate, a maturation score was computed as the Spearman’s rank correlation between the individual gradient and the group template, reflecting the similarity between the individual gradient and those in the average neonate^56^.

The first two gradients, which explained 41.0% and 14.4% of the variance, respectively, were selected. To compare functional gradients in neonates and adults, Spearman’s rank correlation was performed. Adult functional gradient data (cortex-only) were downloaded using the neuromaps toolbox (https://netneurolab.github.io/neuromaps/), originally calculated by Margulies et al.^1^. For between-group comparisons, neonatal functional gradients, originally computed across the whole brain, were restricted to the cortical regions to match the cortex-only adult data. Vertex-level adult functional gradients were first transformed into the Schaefer-400 atlas using the neuromaps toolbox. Spearman’s correlation was then performed with 1000 spin tests^64^ to control for spatial autocorrelation.

### White matter distances

Connection lengths were estimated using white matter distance, the shortest distance through the interior white matter volume, rather than Euclidean or geodesic distance used in previous studies. White matter forms the anatomical pathways linking brain regions, providing the structural basis for functional connections. Therefore, the white matter distances represent the shortest plausible pathway through white matter by which an anatomical connection could link two regions. We calculated these white matter distances between every pair of cortical regions in the parcellation within the dHCP 40-week template as follows:

1. Converting the white matter volume into a triangular mesh, by making each white matter voxel a node in the mesh and forming edges between voxels that shared a vertex. If this graph contained multiple disconnected components, only the largest component was retained for subsequent analyses.
2. For each cortical region of interest (ROI), voxels were similarly represented as a graph. Voxels at the grey–white matter interface were identified as those sharing a vertex with voxels in the retained white matter mesh. For each ROI, shortest-path distances (computed using Dijkstra’s algorithm) were calculated from these interface voxels to all other voxels within the ROI. The interface voxel with the minimum mean shortest-path distance to all ROI voxels was defined as the ROI centroid.
3. Each ROI centroid was added to the white matter graph as a new node and connected to adjacent white matter nodes with which it shared a vertex.
4. Shortest-path distances between all pairs of cortical ROI centroids were computed on the white matter graph, producing a 400-by-400 distance matrix of cortico-cortical white matter distances (Supplementary Figure S14).
5. Subcortical and cerebellar voxels were represented as a separate graph. For each region, the centroid was defined as the voxel minimising the mean shortest-path distance to all other voxels within that region. A combined subcortical/cerebellar graph was constructed, excluding connections between voxels in opposite hemispheres. This graph was then integrated with the white matter graph by connecting subcortical voxels to adjacent white matter voxels that shared a vertex. Shortest-path distances between all subcortical/cerebellar and cortical centroids were then computed. These were combined with cortico-cortical distances to produce a final 433-by-433 matrix of white matter distances between all brain regions.

### Long- and short-range functional connectivity

Connection lengths ranged from 4.10 mm to 122.04 mm, and the mean functional connectivity strength (FCS) decreased with connection length (Supplementary Figure S14). Functional connections were categorised into three bins: short (< 40 mm), middle (40-80 mm), and long-range (> 80 mm) connections.

Proportional thresholding was applied across a range of network densities (5% to 50%, at 1% steps). For each node, normalised functional connectivity strength (normalised FCS) within each distance bin was computed as the sum of strength (i.e., sum of functional connectivity) to all other nodes in that bin divided by the node’s total strength. Normalised FCS reflects the proportion of short-, middle-, and long-range connections for each node, with the sum across the three bins equal to 1. At the network level, normalised FCS was obtained by averaging nodal values within each network.

### Associations between functional gradients and distance-dependent functional connectivity

To examine the correspondence between distance-dependent connectivity and the development of neonatal functional gradients, we computed Spearman’s rank correlations between functional gradient maps and normalised FCS maps at both individual and group levels. At the group level, correlations were calculated between the group-averaged normalised FCS map in all term-born neonates and the group template of functional gradients across the whole brain. To account for spatial autocorrelation, 1000 spin tests were performed for cortical regions only. In addition to whole-brain analyses, system-level and network-level analyses were also conducted in cortical regions only, due to constraints related to network assignment and spin test. The seven cortical sub-networks were divided into primary regions (i.e., visual and sensorimotor networks) and association regions (i.e., default-mode, frontoparietal, dorsal attention, salience/ventral attention, and limbic networks). At the individual level, correlations were computed between individual normalised FCS maps and gradient maps across the whole brain.

### Development of functional gradients and distance-dependent functional connectivity

We examined developmental changes in functional gradients and distance-dependent functional connectivity. For functional gradients, to examine both linear and non-linear developmental changes in maturation score (defined as the similarity of individual functional gradient to the average neonate), generalised additive models (GAM) were fitted using the *mgcv* package in R^65^. GAMs were fitted for the first two gradients separately, with PMA at scan modelled as a smooth term and sex and number of head motion outliers as linear covariates: *Maturation score ∼ s(PMA at scan) + Sex + head motion*. The smooth term for PMA at scan produced a smooth function (or spline) as a linear combination of weighted basis functions, capturing the developmental trajectory of the maturation score. The smooth term was modelled using cubic splines as the basis set, with the maximum basis function complexity (k) set to 3, and smoothing parameters estimated using restricted maximum likelihood (REML). The selected basis function complexity (k = 3) resulted in the lowest Akaike information criterion (AIC) and Bayesian information criterion (BIC) for the maturation score of G1, although AIC and BIC values varied little for both G1 and G2 across k = 3-10 (Table S1). To assess the developmental changes, the full model was compared with a reduced model (*Maturation score ∼ Sex + head motion*). Effect size was quantified as the change in adjusted R^2^ (ΔR_adj_^2^) between the two models, and significance was determined by an analysis of variance (ANOVA) comparing the full and reduced models^66^. In addition, the rate of developmental changes was assessed as the first derivative of the smooth term for PMA at scan, estimated using finite differences. A 95% confidence interval of the first derivative was calculated using the *gratia* package in R. Significant change was identified where the confidence interval did not include zero^67,68^.

For distance-dependent functional connectivity, normalised FCS of short-, middle-, and long-range connections were analysed. Linear regression models were fitted to these measures separately, with PMA at scan as the main predictor, controlling for sex and number of head motion outliers.

To further examine *in utero* and *ex utero* maturation, we decomposed the effect of *PMA at scan* into *GA at birth* and *postnatal age*. For neonatal functional gradients, we fitted linear regression models: *Maturation score (G1_neo_/G2_neo_) ∼ GA at birth + PNA at scan + sex + head motion*. For distance-dependent functional connectivity, we fitted linear regression models: *Normalised FCS (long/middle/short) ∼ GA at birth + PNA at scan + sex + head motion*.

### Connectivity distance

We computed connectivity distance, a continuous measure defined for each node as the average white matter distance to its functionally connected nodes after proportional thresholding of the functional connectivity matrix^21^. To complement the normalised FCS results within each distance bin, connectivity distance was examined using the same analytical framework. At the group level, we computed Spearman’s rank correlations between the group-level functional gradient maps and the group-averaged connectivity distance map across the whole brain, with 1000 spin tests for cortical regions only. System-level and network-level analyses were also conducted in cortical regions only.

### Statistics

We examined the developmental changes in functional gradients using generalised additive models. Statistical significance was assessed using ANOVA comparing the full model (with a smooth term for PMA at scan) with a reduced model without PMA at scan. Effect sizes are reported as the change in adjusted R^2^ between the two models.

We examined the developmental effects of distance-dependent functional connectivity using linear regression models. Statistical significance was assessed using two-sided t-tests. Effect sizes are reported as partial correlation coefficients.

When comparing brain maps (i.e., neonatal and adult functional gradients; distance-dependent functional connectivity and functional gradients), we calculated Spearman’s rank correlations. To account for spatial autocorrelation in cortical regions, 1000 spin tests were performed across the cortical regions to generate a null distribution. The statistical significance was determined by comparing the empirical correlation to the null distribution^64,69^.

For network-level analysis, *p* values were further corrected using the Benjamini-Hochberg false discovery rate (FDR) method^70^. Adjusted *p* values were reported.

## Disclosure of Delegation to Generative AI

The authors declare the use of generative AI in the research process. According to the GAIDeT taxonomy (2025), Code optimisation task was delegated to GAI tools under full human supervision. The GAI tool used was: ChatGPT-5. Responsibility for the final manuscript lies entirely with the authors. GAI tools are not listed as authors and do not bear responsibility for the final outcomes.

## Code and Data Availability

The data used in this study were obtained from the Developing Human Connectome Project (dHCP). dHCP data are available through the dHCP data repository: https://data.developingconnectome.org.

All code that we used is available in https://github.com/QianwenChang/Neonatal_functional_gradient.

## Acknowledgments

This work was supported by the European Research Council under the European Union’s Seventh Framework Programme (FP7/20072013)/ERC grant agreement no. 319456 (dHCP project). This work was also partly funded through philanthropic support from Heart of Racing LLC to the Brain Health Gen2020 programme at King’s College London, and by the National Institute for Health Research (NIHR) Biomedical Research Centre (BRC): Maudsley. The authors also acknowledge support from the Institute for Translational Neurodevelopment at King’s College London, the European Autism Interventions (EU-AIMS) trial and the EU AIMS-2-TRIALS, a European Innovative Medicines Initiative Joint Undertaking under Grant Agreements No. 115300 and 777394, the resources of which are composed of financial contributions from the European Union’s Seventh Framework Programme (Grant FP7/2007–2013) and Horizon 2020 from the European Federation of Pharmaceutical industries and Associations companies’ in-kind contributions. QC received support from KCL-CSC joint scholarship awarded by King’s College London and Chinese Scholarship Council. NT, TA and GM received support from the Medical Research Council Centre for Neurodevelopmental Disorders, King’s College London [MR/N026063/1]. TA was supported by a Medical Research Council (MRC) Senior Clinical Fellowship [MR/Y009665/1]. SO was supported by The Marian and E.H. Flack Trust. The views expressed are those of the authors and not necessarily those of the funders, IHI-JU2, King’s Health Partners, the NHS, the National Institute for Health Research, or the Department of Health and Social Care. The funders had no role in the design and conduct of the study; collection, management, analysis, and interpretation of the data; preparation, review, or approval of the manuscript; and decision to submit the manuscript for publication.

## Author Contributions

Q.C., and D.B. designed and performed the research. Q.C. analysed the data. S.O. contributed new analytical tools. S.F.M. pre-processed data. Q.C. and D.B. wrote the manuscript. Q.C., S.O., S.F.M., N.T., T. A., G.M., and D.B. helped to interpret results, edited and reviewed the manuscript. G.M., and D.B. supervised the study.

## Competing Interest Statement

G.M. has received funding for investigator-initiated studies from GW Pharmaceuticals and COMPASS Pathfinder Ltd., G.M. has consulted for Greenwich Biosciences, Inc. Q.C., S.O., S.F.M., N.T., T.A., and D.B. have no conflicts to declare.

## Notes

### Competing Interest Statement

G.M. has received funding for investigator‐initiated studies from GW Pharmaceuticals and COMPASS Pathfinder Ltd., and has consulted for Greenwich Biosciences, Inc. Q.C., S.O., S.F-M., N.T., T.A., and D.B. have no conflicts to declare.

### Summary of Updates

Added sensitivity analyses and edited causal language.

