## Supplementary Figure for "Emergence of long- and short-range functional connectivity parallels the development of neonatal brain gradients"

### Supplementary Figures

#### 1. Validation analysis in the oldest term-born neonates (23 term-born neonates, PMA at scan ≥ 44 weeks)

##### 1.1 Functional gradients in a subgroup of the oldest term-born neonates

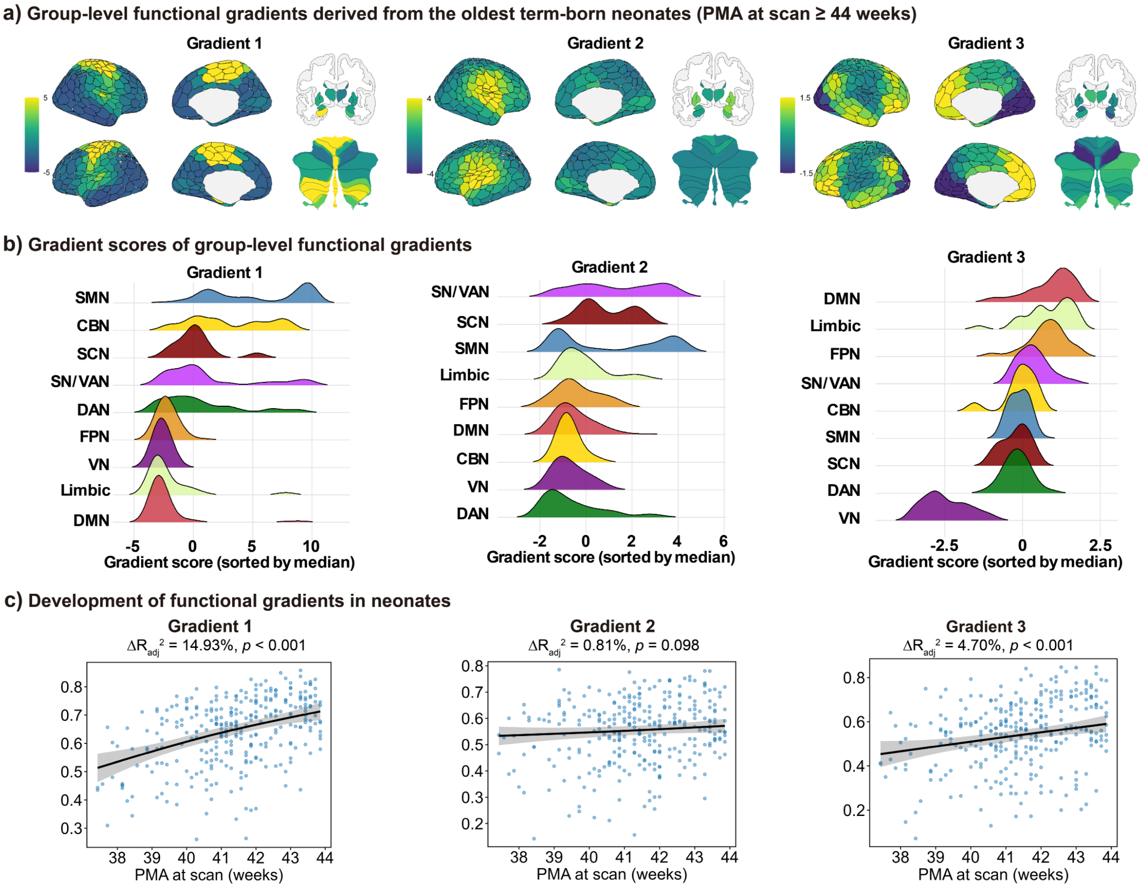


**Fig. S1.** (a) The first three group-level functional gradients in the oldest term-born neonates (23 neonates, PMA at scan ≥ 44 weeks), explaining 41.0%, 14.4%, and 13.9% of the variance, respectively. (b) Gradient scores for each network. Networks are ordered by the median gradient score. SMN = sensorimotor network, CBN = cerebellar network, SCN = subcortical network, SN/VAN = salience/ventral attention network, DAN = dorsal attention network, FPN = frontoparietal network, VN = visual network, DMN = default-mode network. (c) Development of functional gradients. Maturation scores of the first and third gradients increased significantly with PMA at scan, after controlling for sex and in-scanner head motion (FD outliers), modelled using the GAM. Effect size and significance were determined by comparing the full model with a reduced model excluding the smooth term of PMA at scan.

#### 1.2 Correspondence between functional gradients in adults and the oldest term-born neonates

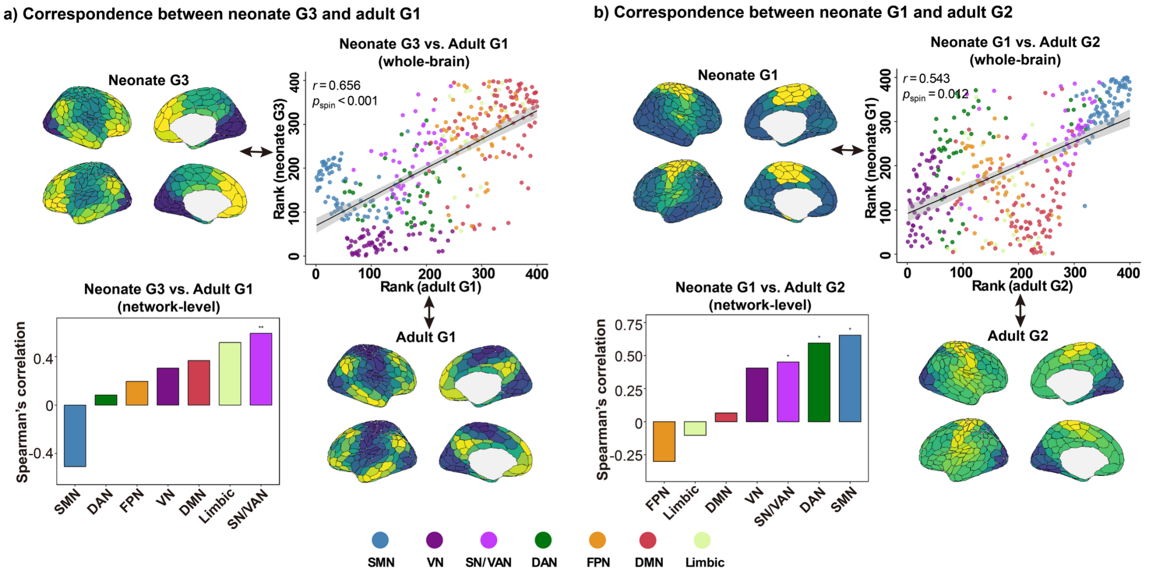


**Fig. S2.** (a) Correspondence between the third gradient (G3) in the oldest term-born neonates and the primary gradient (G1) in adults at the whole-brain and network levels. (b) Correspondence between the first gradient (G1) in the oldest term-born neonates and the secondary gradient (G2) in adults at the whole-brain and network levels. Spearman's rank correlations were calculated between the two brain maps, with 1000 spin tests performed for both whole-brain and network-level analyses. $^{**}p_{\text{spin}} < 0.01$, $^{*}p_{\text{spin}} < 0.05$.

##### 1.3 Associations between distance-dependent functional connectivity and functional gradients in the oldest term-born neonates

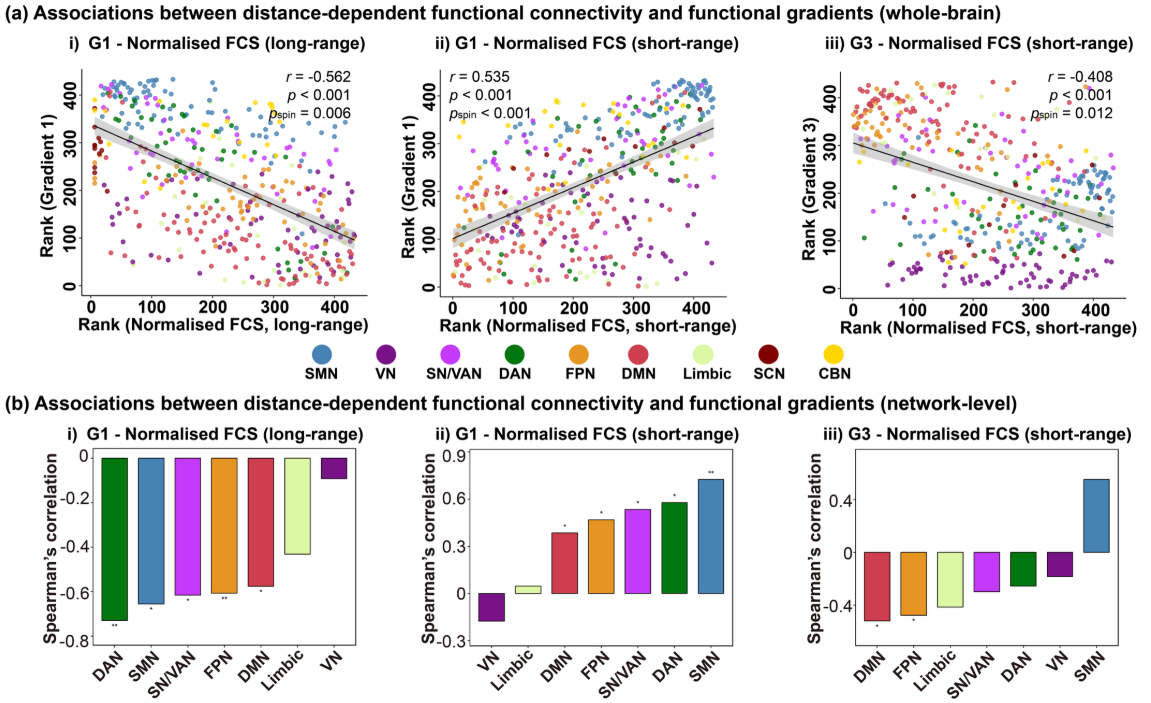


**Fig. S3.** Associations between group-averaged distance-dependent functional connectivity and group templates of functional gradients in the oldest term-born neonates at the (a) whole-brain and (b) network levels. Spearman's rank correlations were calculated between the group-averaged normalised FCS and the group templates of functional gradient maps with 1000 spin tests. The G1 template showed opposing associations with long- and short-range functional connections. The G3 template was negatively associated with short-range connections. SN/VAN = salience/ventral attention network; SMN = sensorimotor network; DMN = default-mode network; DAN = dorsal attention network; FPN = frontoparietal network; VN = visual network. $^{**}p_{spin} < 0.01$, $^{*}p_{spin} < 0.05$.

##### 1.4 Associations between connectivity distance and functional gradients in the oldest term-born neonates

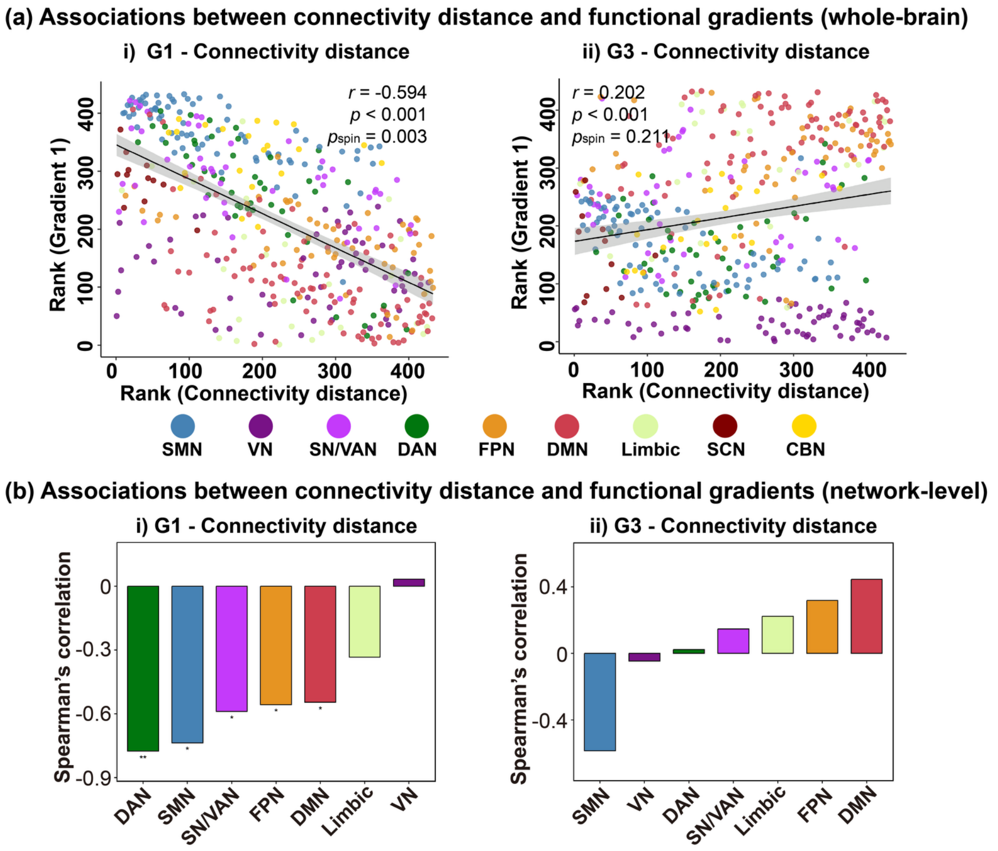


**Fig. S4.** Associations between group-averaged connectivity distance and group templates of functional gradients in the oldest term-born neonates at the (a) whole-brain and (b) network levels. Spearman's rank correlations were calculated with 1000 spin tests. The G1 template was negatively associated with the CD map, while no significant association was found between the G2 template and the CD map across the whole brain. SN/VAN = salience/ventral attention network; SMN = sensorimotor network; DMN = default-mode network; DAN = dorsal attention network; FPN = frontoparietal network; VN = visual network; SCN = subcortical network; CBN = cerebellar network. $^{**}p_{spin} < 0.01$, $^{*}p_{spin} < 0.05$.

#### 2 Correspondence between adult and neonatal functional gradients at the system-level

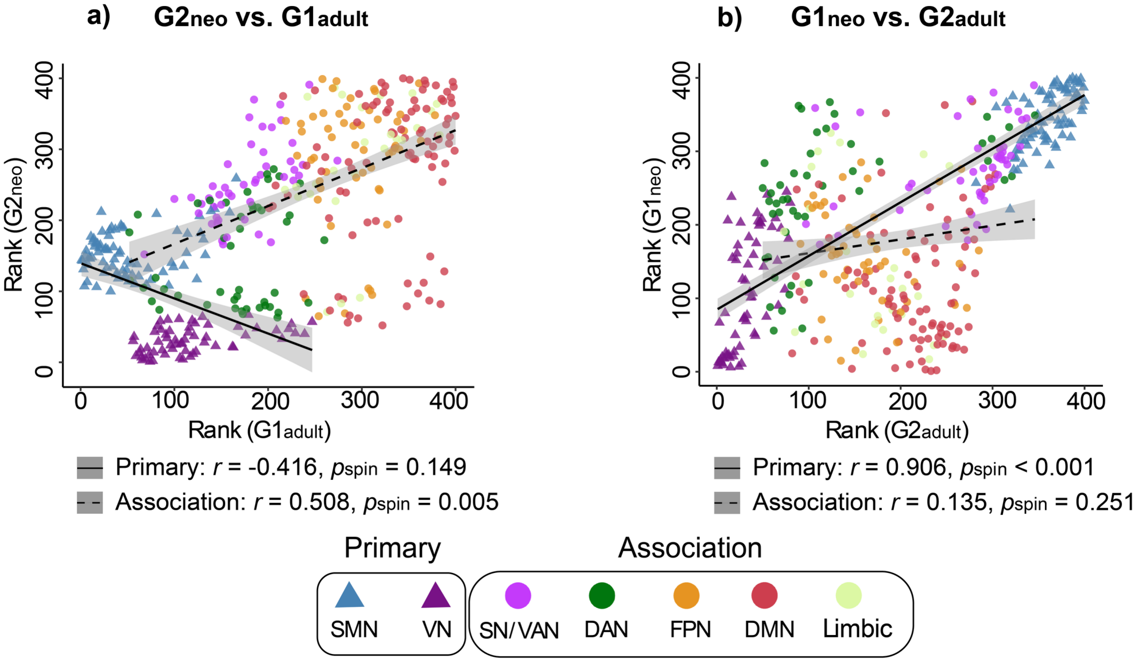


**Fig. S5.** (a) Correspondence between the second gradient in neonates ($G2_{neo}$) and the principal gradient in adults ($G1_{adult}$) at the system level (primary and association regions). (b) Correspondence between the first gradient in neonates ($G1_{neo}$) and the secondary gradient in adults ($G2_{adult}$) at the system level. Spearman's rank correlations were calculated between the two brain maps within each part, with 1000 spin tests.

##### 3 Neonatal functional gradients derived from cortico-cortical functional connectivity

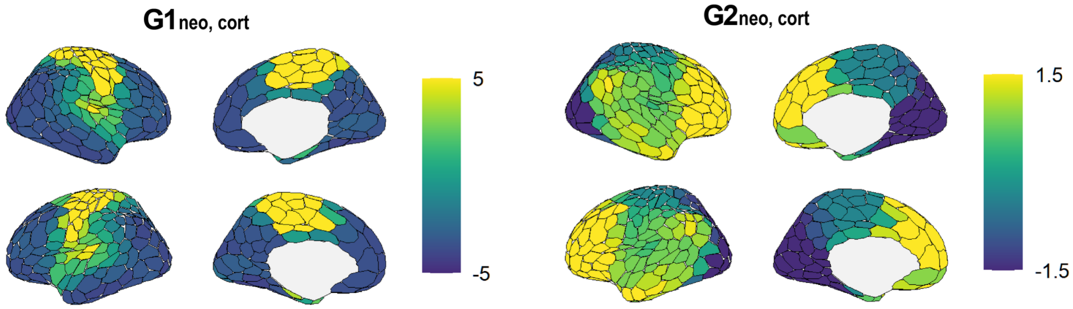


**Fig. S6.** The first two group-level functional gradients ($G1_{neo,cort}$, $G2_{neo,cort}$) were derived from cortico-cortical functional connectivity, explaining 43.5% and 14.8% of the variance, respectively.

#### 4 Development of distance-dependent functional connectivity after controlling for mean FD and mean DVARS (in addition to % FD outliers)

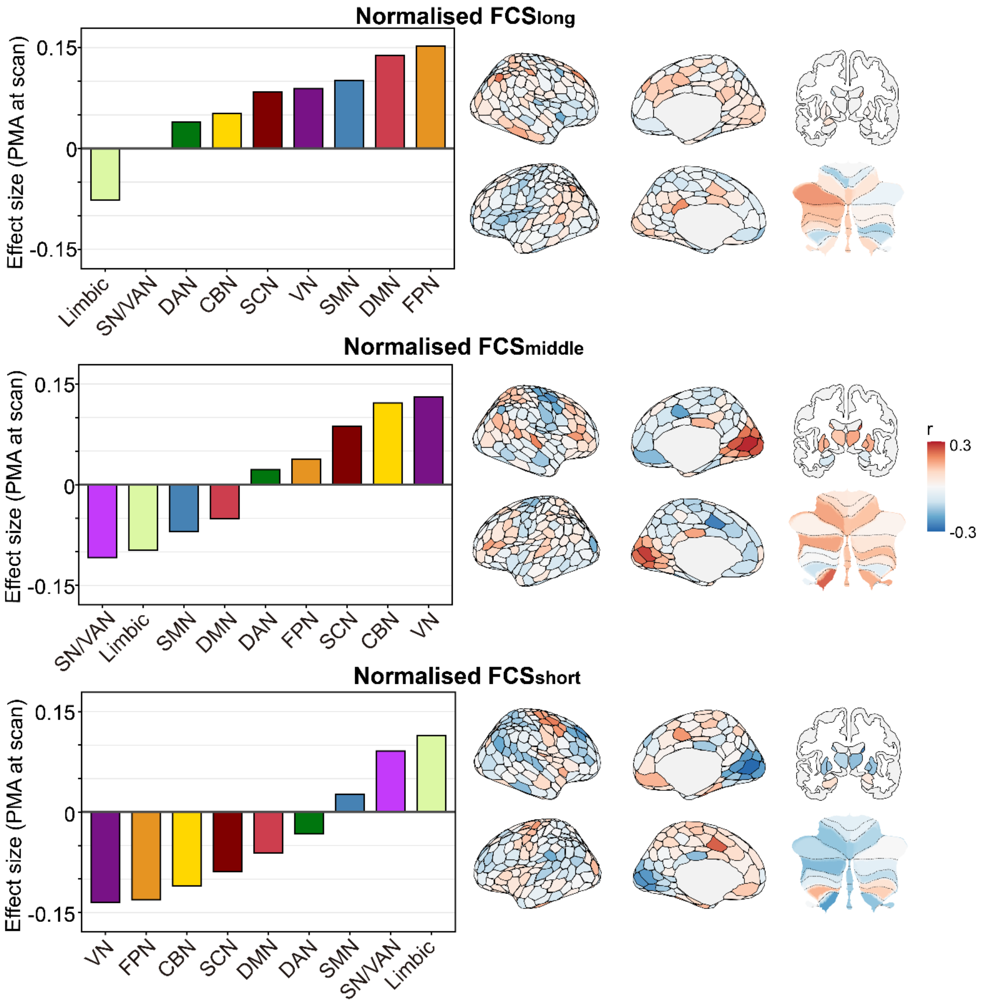


**Fig. S7.** (a) Development of normalised FCS for long-, middle-, and short-range connections at the network (left) and parcel levels (right) when controlling % FD outliers, mean FD, and mean DVARS. Linear regression models were fitted, with PMA at scan as the main predictor, controlling for sex and in-scanner head motion (% FD outliers, mean FD, and mean DVARS). The effect size was reported as the partial correlation coefficient. SN/VAN = salience/ventral attention network; SMN = sensorimotor network; DMN = default-mode network; DAN = dorsal attention network; FPN = frontoparietal network; VN = visual network; SCN = subcortical network; CBN = cerebellar network. $^{*}p_{FDR} < 0.05$, FDR corrected across 9 networks.

#### 5 *In utero* and *ex utero* maturation of distance-dependent functional connectivity

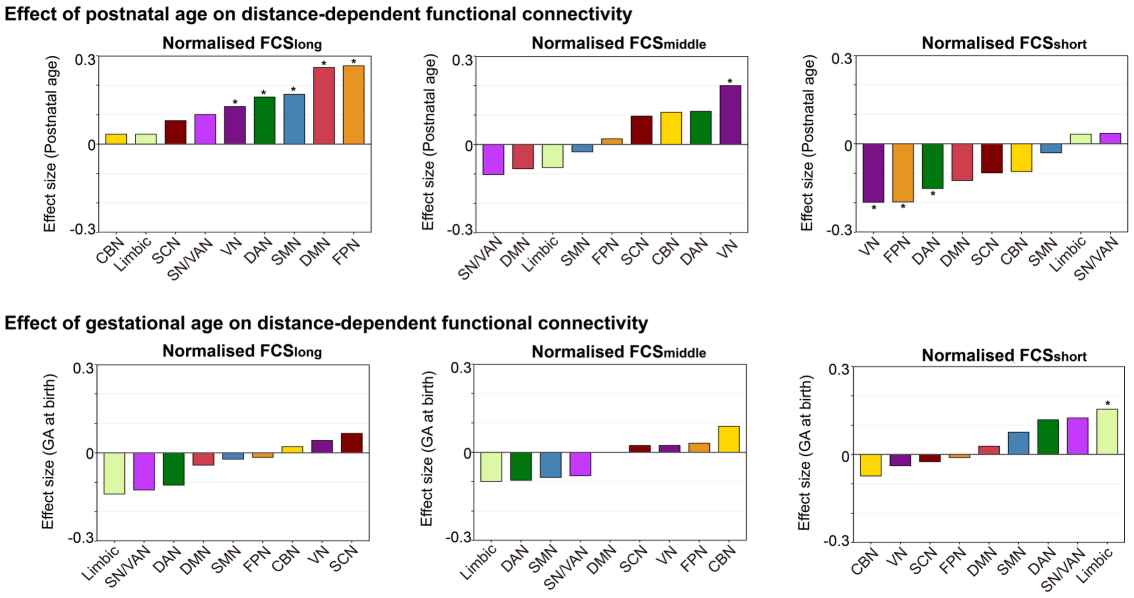


**Fig. S8.** *In utero* and *ex utero* maturation of distance-dependent functional connectivity. Effects of postnatal age (PNA) at scan (top) and gestational age (GA) at birth (bottom) were estimated using linear regression models that included PNA at scan and GA at birth as separate terms and controlled for sex and in-scanner head motion (FD outliers). Effect sizes are reported as partial correlation coefficients. *$p_{FDR}$ values < 0.05, FDR corrected across 9 networks. SMN = sensorimotor network, CBN = cerebellar network, SCN = subcortical network, SN/VAN = salience/ventral attention network, DAN = dorsal attention network, FPN = frontoparietal network, VN = visual network, DMN = default-mode network.

#### 6 Developmental changes in associations between distance-dependent functional connectivity and functional gradients

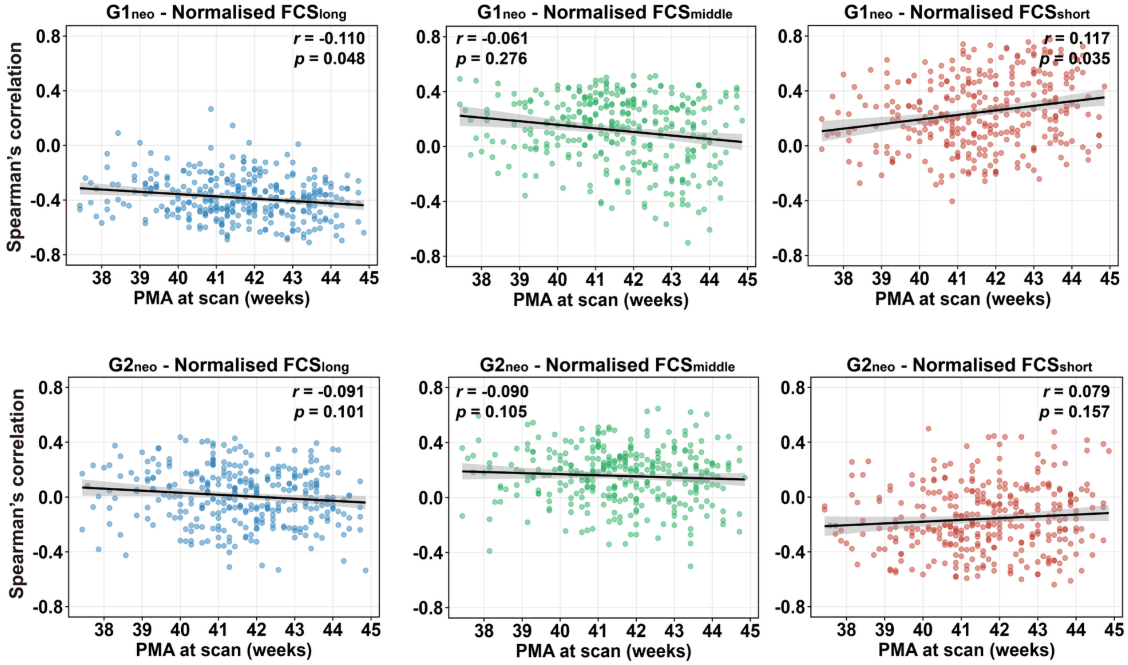


**Fig. S9.** Developmental changes in associations between distance-dependent functional connectivity and functional gradients. Each dot represents Spearman's rank correlation value between functional gradient map ($G1_{neo}$, $G2_{neo}$) and normalised FCS map (normalised $FCS_{long}$, normalised $FCS_{middle}$, and normalised $FCS_{short}$) for each neonate. Linear regression was fitted to the correlation values, with PMA at scan as the main predictor, controlling for sex and in-scanner head motion (FD outliers). The effect size was reported as the partial correlation coefficient. *P*-values were uncorrected.

#### 7 Split-half analyses

##### 7.1 Function gradients estimated from the first half of the resting-state data timeseries (half 1) in all term-born neonates

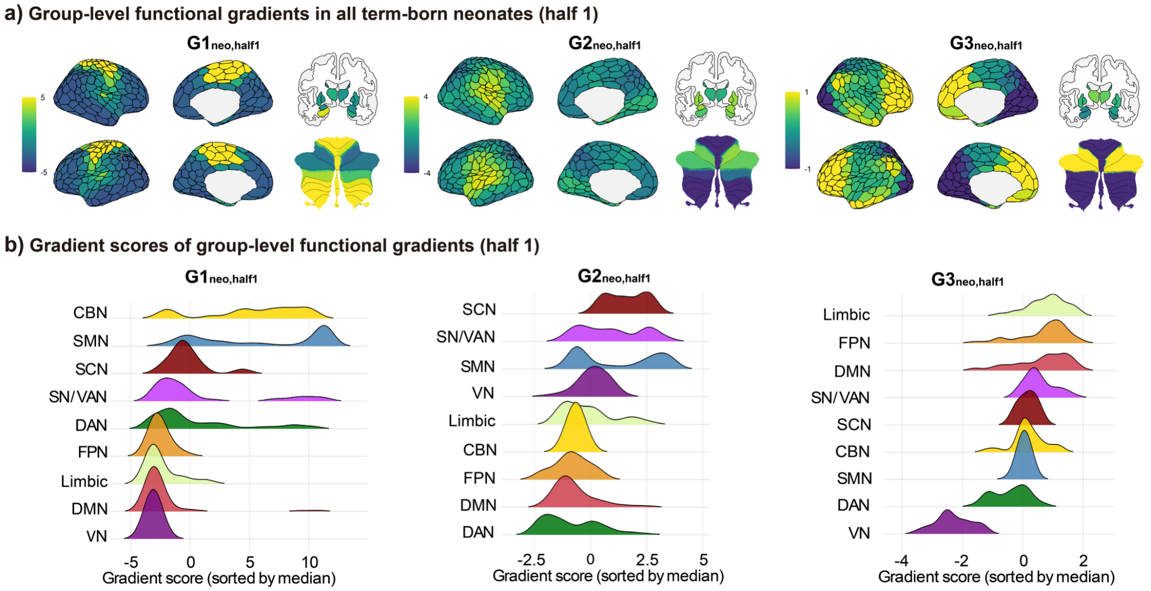


**Fig. S10.** Functional gradients estimated from the first half of the resting-state data timeseries in all term-born neonates. (a) The first three group-level functional gradients in all term-born neonates, explaining 46.1%, 14.0%, and 11.5% of the variance, respectively. (b) Gradient scores for each network. Networks are ordered by the median gradient score. SMN = sensorimotor network, CBN = cerebellar network, SCN = subcortical network, SN/VAN = salience/ventral attention network, DAN = dorsal attention network, FPN = frontoparietal network, VN = visual network, DMN = default-mode network**.**

##### 7.2 Development of distance-dependent functional connectivity estimated from the second half of the resting-state data timeseries (half 2) in all term-born neonates

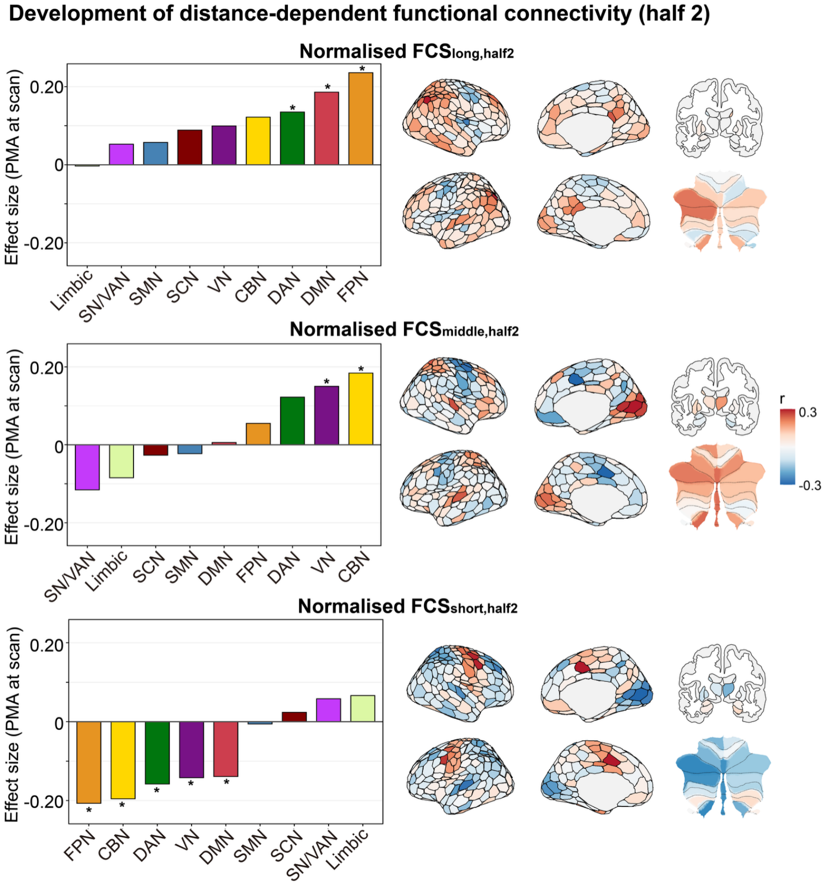


**Fig. S11.** (a) Development of normalised FCS for long-, middle-, and short-range connections, estimated from the second half of each neonate's resting-state data timeseries, at the network (left) and parcel levels (right). Linear regression models were fitted, with PMA at scan as the main predictor, controlling for sex and in-scanner head motion (FD outliers). The effect size was reported as the partial correlation coefficient. SN/VAN = salience/ventral attention network; SMN = sensorimotor network; DMN = default-mode network; DAN = dorsal attention network; FPN = frontoparietal network; VN = visual network; SCN = subcortical network; CBN = cerebellar network. *$p_{FDR}$ < 0.05, FDR corrected across 9 networks.

#### 7.3 Associations between distance-dependent functional connectivity (half 2) and functional gradients (half 1)

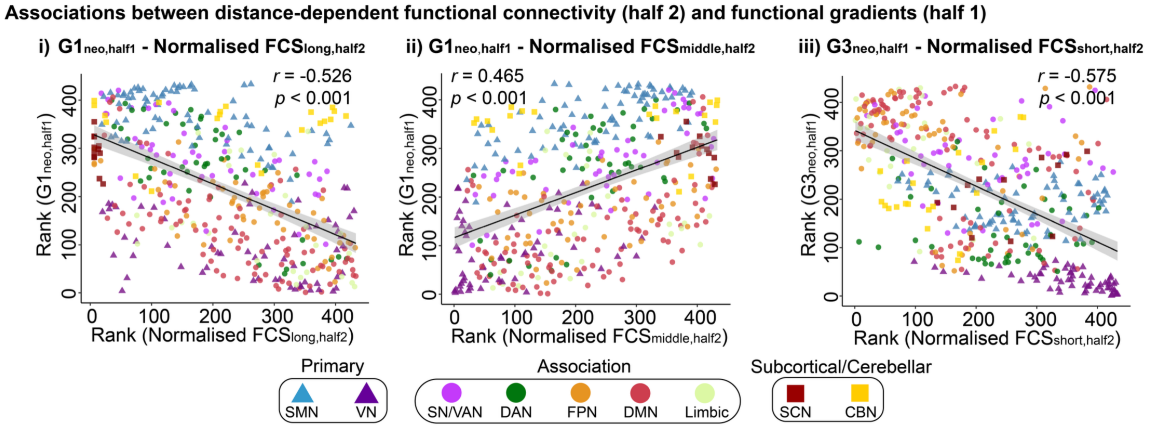


**Fig. S12.** Associations between group-averaged distance-dependent functional connectivity (half 2) and functional gradients (half 1) across the whole brain. Spearman's rank correlations were calculated between group-averaged normalised FCS and functional gradient maps. $G1_{neo,half1}$ showed opposing associations with normalised $FCS_{long,half2}$ ($r$ = -0.526, $p$ < 0.001; with 1000 spin tests in cortical regions, $r$ = -0.564, $p_{spin}$ = 0.006) and normalised $FCS_{middle,half2}$ ($r$ = 0.465 $p$ < 0.001; with 1000 spin tests in cortical regions, $r$ = 0.480, $p_{spin}$ = 0.021). $G3_{neo,half1}$ was negatively associated with normalised $FCS_{short,half2}$ ($r$ = -0.575, $p$ < 0.001; with 1000 spin tests in cortical regions, $r$ = -0.611, $p_{spin}$ = 0.002). No significant associations were found for the remaining combinations ($p_{spin}$ values > 0.05).

#### 7.4 Associations between connectivity distance (half 2) and functional gradients (half 1)

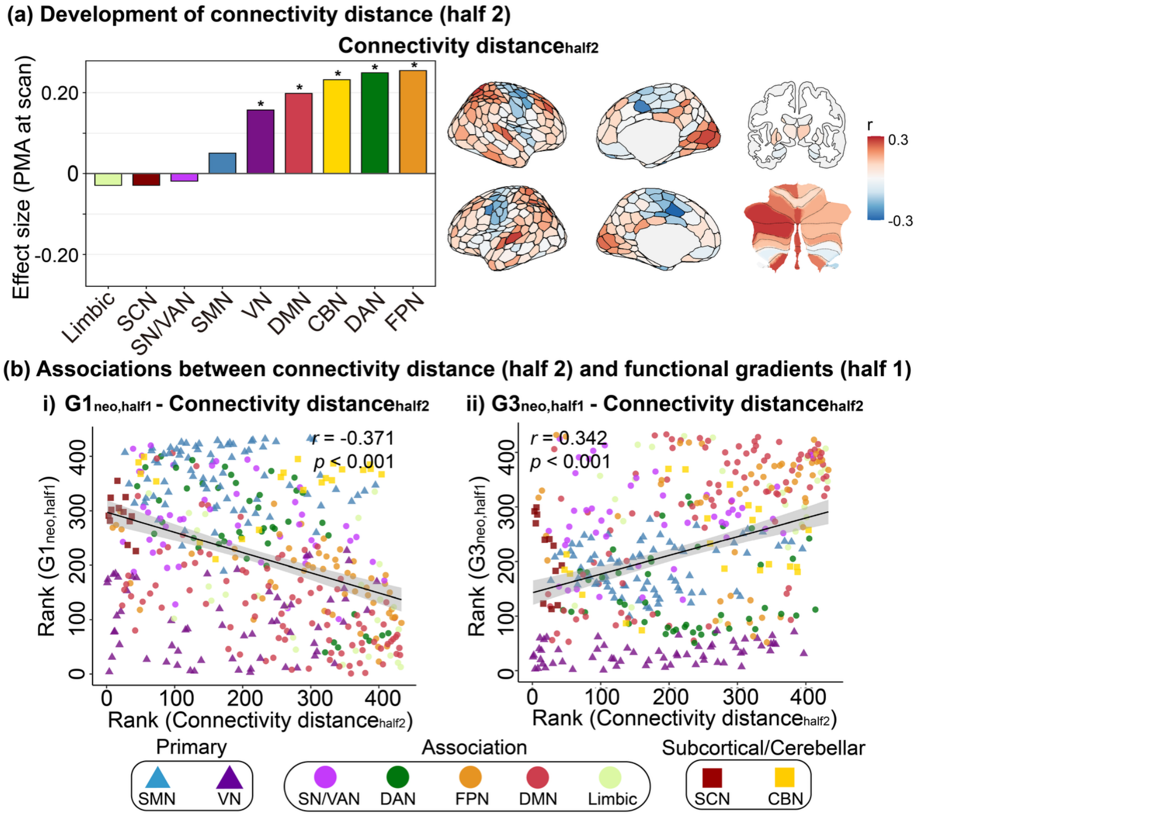


**Fig. S13.** (a) Development of connectivity distance, estimated from the second half of each neonate's resting-state data timeseries, at the network (left) and parcel levels (right). Linear regression models were fitted, with PMA at scan as the main predictor, controlling for sex and in-scanner head motion (FD outliers). The effect size was reported as the partial correlation coefficient. $^{*}p_{FDR} < 0.05$, FDR corrected across 9 networks. (b) Associations between group-averaged connectivity distance (half 2) and functional gradients (half 1) across the whole brain. Spearman's rank correlations were calculated. $G1_{neo,half1}$ was negatively associated with the $CD_{half2}$ map ($r$ = -0.371, $p < 0.001$; with 1000 spin tests in cortical regions, $r$ = -0.400, $p_{spin}$ = 0.025), while $G3_{neo,half1}$ was positively associated with the $CD_{half2}$ map across the whole brain ($r$ = 0.342, $p < 0.001$; with 1000 spin tests in cortical regions, $r$ = 0.355, $p_{spin}$ = 0.010). SN/VAN = salience/ventral attention network; SMN = sensorimotor network; DMN = default-mode network; DAN = dorsal attention network; FPN = frontoparietal network; VN = visual network; SCN = subcortical network; CBN = cerebellar network.

#### 8 Connection length estimated by white matter distance

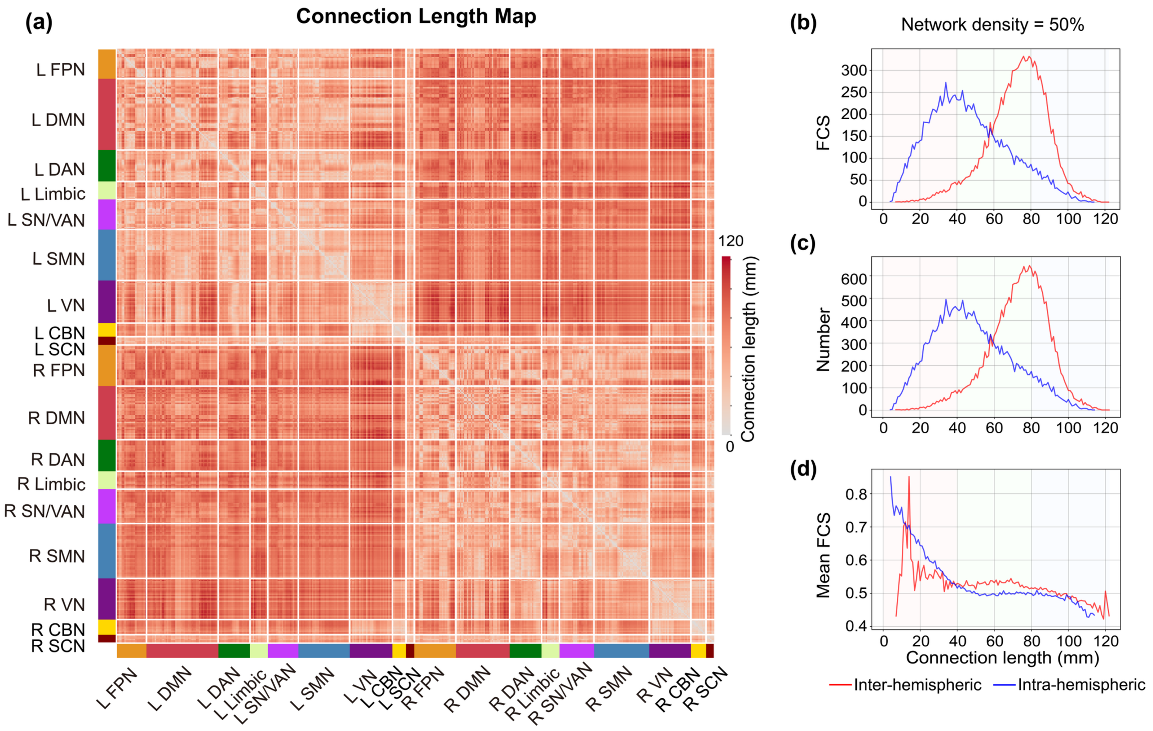


**Fig. S14.** (a) Connection length map estimated by white matter distance. L = left hemisphere, R = right hemisphere. (b) sum of functional connectivity strength (FCS); (c) the number and (d) mean functional connectivity strength (mean FCS) across different connection lengths at the network density of 50%. Red line denotes inter-hemispheric connections; blue line denotes intra-hemispheric connections. The number and FCS of inter-hemispheric connections peaked around 80 mm, while the number and FCS of intra-hemispheric connections peaked around 40 mm, the intersection of the two lines was at around 60 mm. The mean FCS of functional connections decreased with connection length.

#### Supplementary Tables

**Table S1.** AIC and BIC values of the GAM models across k ranging from 3 to 10

| Basis function complexity (k) | AIC | BIC |
|---|---|---|
| **G1** | | |
| 3 | -517.5502 | -494.9632 |
| 4 | -516.9657 | -493.2296 |
| 5 | -517.1478 | -492.6002 |
| 6 | -516.9098 | -492.2605 |
| 7 | -516.9300 | -492.1134 |
| 8 | -517.1343 | -489.3717 |
| 9 | -516.9006 | -491.9607 |
| 10 | -517.1011 | -489.1702 |
| **G2** | | |
| 3 | -322.8710 | -301.4095 |
| 4 | -323.1299 | -302.5012 |
| 5 | -323.0867 | -302.4459 |
| 6 | -323.0659 | -302.4419 |
| 7 | -323.0648 | -302.4524 |
| 8 | -322.4737 | -299.6768 |
| 9 | -323.0571 | -302.4718 |
| 10 | -322.4452 | -299.5617 |
